# MPNN-guided redesign of PET hydrolases with enhanced catalytic activity below the PET glass transition temperature

**DOI:** 10.64898/2026.02.25.708052

**Authors:** Aransa Griñen, Valeria Eltit-Villarroel, Francisca Durán-Osorio, Javiera Avilés, Flavia C. Zacconi, Edson N. Cárcamo Noriega, Christopher D. Bahl, Ben A. Meinen, César A. Ramírez-Sarmiento

## Abstract

Enzymatic depolymerization of polyethylene terephthalate (PET) presents a sustainable route for plastic circularity, but its industrial viability is disadvantaged by the need for thermostable enzymes active under mild, energy-efficient conditions. While Polyester Hydrolase Leipzig 7 (PHL7, also known as PES-H1) rapidly degrades amorphous PET near the glass transition temperature of this polymer (∼65°C), its poor protein expression, inactivation above 60°C and slow depolymerization below 60°C limits its practical application. Here, we employ ProteinMPNN and LigandMPNN, structural and evolutionary information, to redesign the sequence of PHL7 and improve protein expression, thermostability and activity. We identified 2/36 experimentally tested variants (D5, D11) with enhanced PET depolymerization at 50°C, achieving the same efficiency as PHL7 at 70°C but with a shifted product profile, favoring mono-(2-hydroxyethyl) terephthalate (MHET) over terephthalate. Molecular dynamics revealed that these redesigns exhibit enhanced flexibility in active site regions, providing a mechanistic understanding of their low-temperature catalysis. These variants enable a potential route to resynthesize virgin PET via MHET polycondensation, offering an efficient circular economy pathway.

## INTRODUCTION

Polyethylene terephthalate (PET) is one of the most widely used plastics worldwide, with approximately 26.7 million tonnes produced globally in 2024^1^. Most PET-based products, such as bottles and food packaging, are classified as single-use plastics, designed for short-term use and rapid disposal, making PET one of the most discarded polymers and a major source of environmental pollution^2,3^.

To address this waste problem, enzymatic degradation of PET has emerged as a promising complement to conventional recycling^4^, particularly on additional plastic feedstocks that are excluded from mechanical recycling, such as textiles, fibers, layered packaging materials^5^ and thermoformed PET^6^. PET-degrading enzymes, also termed PET hydrolases (PETases), operate under mild, aqueous conditions, without requiring additional chemicals, thereby minimizing the formation of side products^7,8^. Their high specificity allows them to depolymerize PET into high-purity terephthalic acid (TPA) and ethylene glycol (EG), as well as the intermediate mono-(2-hydroxyethyl) terephthalate (MHET). TPA and EG can then be repolymerized into new, virgin-quality PET in a closed-loop recycling process^6,9,10^.

Nevertheless, PETases retrieved from natural environments are not optimized for their use in industrial recycling processes, which prevents their large-scale application^11^. Characteristics such as thermostability, activity, and low protein expression are problematic features in most of the currently discovered natural PET-degrading enzymes, making them key bottlenecks for economic viability at industrial scale^5^.

One of the latest natural enzymes discovered that exhibits some of these desirable characteristics is Polyester Hydrolase Leipzig 7 (PHL7; also known as PES-H1^12^; EC 3.1.1.74; GenBank accession ID SAY37592.1), a methagenomic enzyme from *Thermoanaerobacter* sp. (TaxID: 399726) that has a melting temperature (T_m_) of 79°C and can depolymerize amorphous PET film after 16 h of incubation at around 70°C^6^. Despite this, its application is limited by its low protein yield upon recombinant expression in *Escherichia coli* and its work at temperatures close to its Tm^13^. Given its potential applications in industrial biorecycling processes, these drawbacks translate into a fast reduction in its activity (potentially due to inactivation^14^) and a higher enzyme production cost on a large scale^5^.

The design of enzymes, including PET hydrolases, remains a difficult task. Enzymes are highly specific, and small modifications via residue substitutions can inadvertently cause significant destabilization and activity losses^15,16^. Intending to improve key features of PHL7 without losing its PET depolymerization efficiency, we used deep learning fixed-backbone sequence design tools ProteinMPNN^17^ and LigandMPNN^18^ in combination with evolutionary information, which have been previously demonstrated to allow optimization of the protein yield, thermal and long-term stability and activity of several enzymes^19^. Our strategy focused on the computational sequence redesign of PHL7 and its subsequent screening to facilitate its industrial implementation. Serendipitously, our sequence redesign strategy led to enzyme variants with enhanced catalytic performance below the glass transition temperature of PET, conditions in which PET hydrolysis is typically compromised.

## RESULTS

To broadly explore sequence space constrained by protein structure, we used inverse folding to generate a diverse library of variants. We employed a fixed-backbone sequence redesign strategy using ProteinMPNN^17^ and LigandMPNN^18^ to enhance the expression, stability and activity of PHL7. Our approach followed the methodology established by Sumida et al.^19^, which integrates evolutionary information with ProteinMPNN, and further expanded it with the use of LigandMPNN.

We used the crystallographic structure of TPA-bound PHL7 (PDB ID 8BRB) as the input structure. Active site residues were defined as those within 6 Å from the bound ligand (Figure 1A). Following Sumida et al.^19^, fixing only active site residues during ProteinMPNN or LigandMPNN redesign is insufficient to maintain enzyme activity. Evolutionary information derived from multiple sequence alignments (MSAs) must also guide the selection of conserved residues to fix during redesign. We used Uniref30^20^ with hhblits^21^ to generate 16 MSAs (see Methods) to rank amino acids at each position by their sequence conservation at 30%, 50%, and 70% cutoffs.

**Figure 1.**
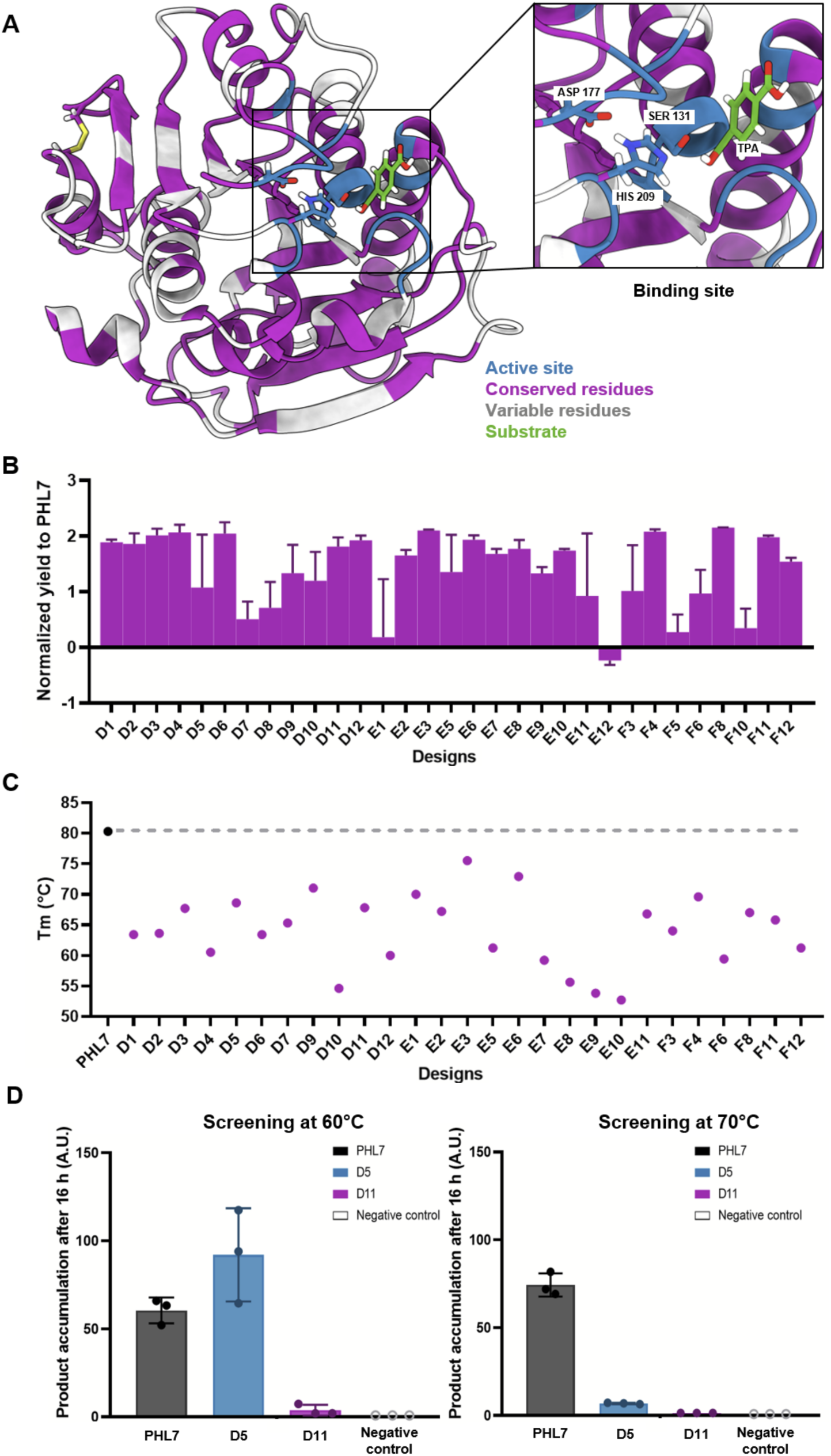
Analysis and screening of redesigned PHL7 variants using deep learning tools. (A) Cartoon representation of TPA-bound PHL7 (PDB ID: 8BRB), with active site residues that are fixed during sequence redesign highlighted in blue, and residues at 50% sequence conservation based on evolutionary information that are fixed during sequence redesign highlighted in purple. The bound TPA in the active site is shown as green sticks, and was used to select the active site residues at a radius of 6 Å. (B) Production of sequence redesigned PHL7 variants measured as normalized yield compared to PHL7, in logarithmic scale. The purification was performed in 2 mL and in duplicates. (C) Melting temperature (Tm) of sequence redesigned PHL7 variants (in °C). (D) PET degradation by active PHL7 redesigns D5 and D11, measured as the accumulation of soluble products after incubation of post-consumer amorphous PET films (half-circle, diameter of 6 mm) for 16 h with the enzymes (9.05 µg/mL) by absorbance at 240 nm. Assays were performed in triplicates at 60 and 70°C. A reaction without enzyme was used as a negative control.

From the 16 generated MSAs, 49 amino acid groups satisfying these constraints were selected and fixed during ProteinMPNN and LigandMPNN sequence redesign (Table S1), generating 1176 sequences (588 from each MPNN model). Three-dimensional structures were predicted using ESMFold^22^ and filtered by pLDDT (a structural confidence metric) and Cα root-mean-square deviation (RMSD) relative to the PHL7 input structure. Stringent filtering (pLDDT > 85, Cα-RMSD < 1.5 Å) followed by RosettaFastRelax^23^ energy minimization (discarding structures with total energy >-200 kcal/mol) did not reduce the sequence pool. Therefore, sequences were clustered into 36 groups using a Levenshtein distance matrix, prioritizing sequence diversity, for further analysis (Table S2). The average sequence identity of the designed sequences was 77.96% (min = 68.73%, max = 96.91%).

Selected sequences (Table S2) were synthesized as codon-optimized genes for expression in *E. coli* and subjected to high-throughput purification in micro-scale from 2 mL cell cultures (Figure S1). PHL7 and 31 out of 36 redesigned enzymes were successfully expressed and purified (Figure 1B and Figure S1). Most redesigns showed higher yields than PHL7 (Figure 1B), with some exceeding the native enzyme by over 120-fold (purified PHL7 average yield = 5 mg/L culture, purified design average yield = 275 mg/L culture; in 2mL cell culture purification), and only one design (E12) exhibited lower yield than PHL7.

To assess global secondary structure content and thermostability, circular dichroism (CD) and differential scanning fluorimetry (DSF) measurements were performed on the redesigned PHL7 variants. Although 31 variants were successfully expressed, secondary structure and T_m_ could only be reliably determined for 27 designs. CD at 25°C confirmed that most proteins are well-folded (Figure S2). All 27 variants showed decreased T_m_ compared to PHL7 by DSF (Figure 1C and Figure S3), with an average T_m_ of 64°C, a 16°C decrease from PHL7. This is in contrast with the increased thermostability observed for other enzymes in previous works using similar redesign protocols^19^.

Activity was screened by incubating post-consumer amorphous PET films with each enzyme for 16 h. Degradation products were quantified by absorbance at 240 nm (bis-(2-hydroxyethyl) terephthalate [BHET], MHET, and TPA) and by the color change of the pH indicator dye phenol red (red to yellow) at 430 nm due to solution acidification after product release. When tested at 60, 70, and 80°C, only two designs—D5 and D11—showed activity against PET (Figure 1D and Figure S4). At 60°C, D5 activity exceeded that of PHL7 (Figure 1D). No degradation was observed at 80°C, possibly due to PET recrystallization as reported before^14^, but more likely due to enzyme denaturation given their thermal stability.

Designs D5 and D11 showed 28-fold and 66-fold increases in protein yield compared to PHL7 in micro-scale purifications (Figure 1B). To confirm scalability, production was scaled from 2 mL to 1 L cell cultures, with changes in induction (see Methods). D5 and D11 exhibited 7-fold and 13-fold protein yield increases, confirming improved recombinant expression in *E. coli* (Figure S5). Analysis of D5 and D11 revealed decreased thermostability compared to PHL7 in micro-scale purified enzymes after differential scanning fluorimetry (DSF) measurements using GloMelt™ (Figure S3). DSF experiments with SYPRO Orange on 1 L-scale purified proteins confirmed T_m_ decreases of ∼10°C (D5) and ∼12°C (D11) relative to PHL7 in high phosphate concentration buffer (Table 1 and Figure S6).

**Table 1.** Melting temperature for PHL7 and active PHL7 redesigns.

| Enzyme | $T_m$ (°C) |
| --- | --- |
| PHL7 | 82.6 ± 0.2 |
| D5 | 72.8 ± 1.2 |
| D11 | 70.7 ± 0.5 |

The combined DSF results show that all successfully analyzed variants exhibit decreased thermal stability compared to PHL7, and that only 2/31 designs showed PET-degrading activity, albeit at lower temperatures. These findings suggest that mutations in D5 and D11 shifted activity towards lower temperatures than PHL7, compensating for their reduced thermal stability.

To explore activity-stability tradeoffs and optimized experimental conditions, we measured polycaprolactone (PCL) nanoparticle degradation at varying temperatures. Turbidity decrease at 600 nm was monitored upon enzyme addition. The PCL nanoparticles exhibited a particle size of 304 nm, polydispersity index of 0.050, and zeta potential of-40 mV by dynamic light scattering (DLS) and laser doppler anemometry (LDA), respectively.

At 25 µg (19.2 mg_ENZYME_/g_PCL_), both designs showed optimal polyesterase activity near 50°C (Figure S7). After 10 min incubation at 50°C, D5 and PHL7 degraded PCL nanoparticles slightly better than D11. At 65°C, D5 and D11 activity decreased dramatically (∼30% and ∼35%, respectively), while PHL7 showed a smaller decrease (∼15%) between 50 and 65°C (Figure 2).

**Figure 2.**
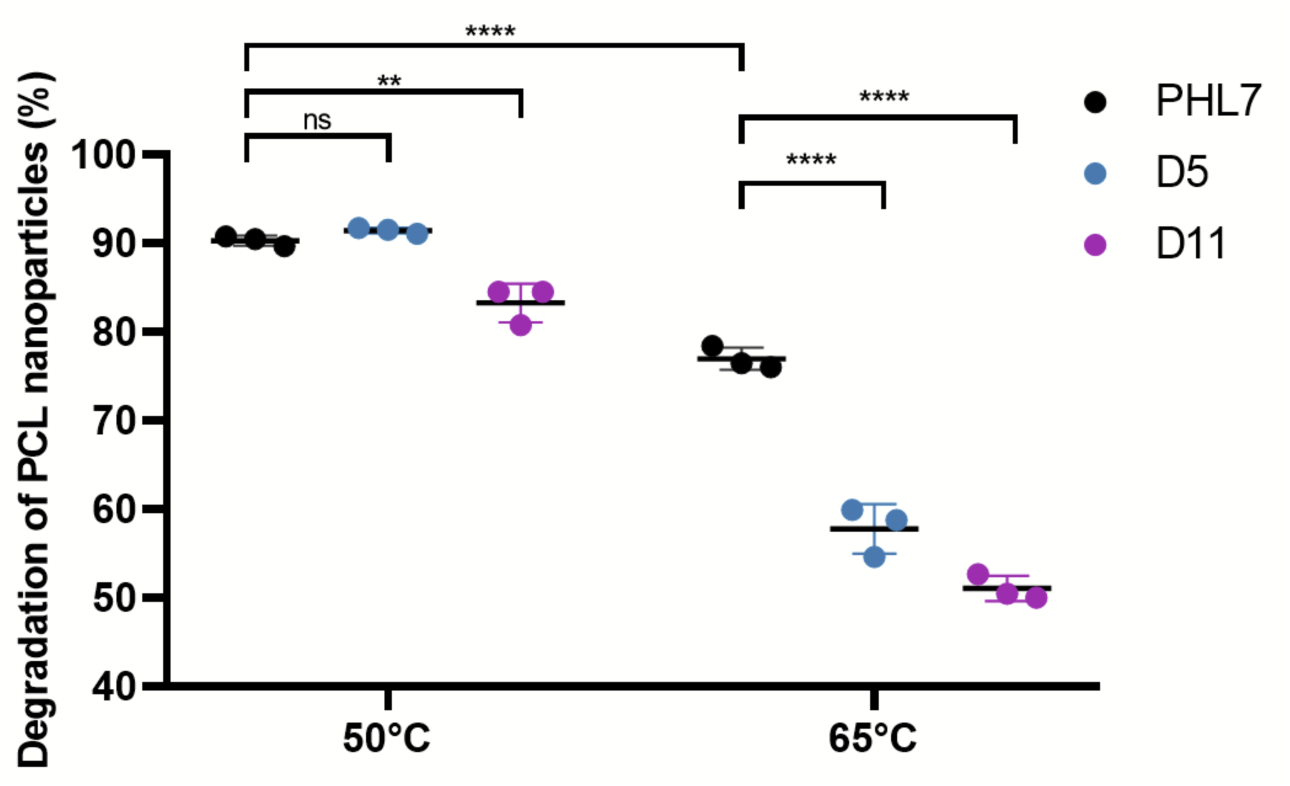
Hydrolysis of PCL nanoparticles by PHL7 and active PHL7 redesigns. Endpoint measurements of PCL nanoparticle degradation at 50°C and 65°C of PHL7, D5 and D11. Assays were performed for 10 mins in triplicates for all enzymes.

Kinetic parameters derived from initial velocity measurements at varying enzyme concentrations (Figure S8)^24^ confirmed that PHL7 exhibits superior kinetics at 50°C compared to D5 and D11 (Table 2). PHL7 showed up to 16-fold faster hydrolysis and up to 4-fold higher affinity for PCL nanoparticles.

**Table 2.** Kinetic parameters of PCL hydrolysis by PHL7 and designs D5 and D11 at 50°C. *k_τ_* is the hydrolysis rate constant and K_A_ as the adsorption equilibrium constant.

| Enzyme | $k_r$ ($10^{-3}/\text{min}$) | $K_A$ (mL/mg) |
| --- | --- | --- |
| PHL7 | 81.24 | 40.8 |
| D5 | 14.94 | 123.0 |
| D11 | 5.16 | 283.2 |

However, PHL7 degrades less post-consumer amorphous PET than D5 after 16 h at 60°C (Figure 1D). This could suggest that PHL7 initially hydrolyzes PET faster but inactivates over time, resulting in lower total degradation. In contrast, D5 exhibits slower rates and lower substrate affinity than PHL7 but apparently maintains activity for longer, achieving similar or superior PET degradation below 70°C.

To test this hypothesis, PET hydrolysis was analyzed. Optimal temperatures were determined by quantifying soluble products (TPA, MHET, BHET) from the degradation of amorphous PET films at different temperatures by absorbance at 240 nm. A 30 min temperature ramp showed optimal performance at 70°C for D5 and 60°C for D11 (Figure S9). Based on these results and the PCL degradation analysis, PET hydrolysis by D5 and D11 was measured at 50, 60, and 70°C.

TPA, MHET and BHET were quantified by HPLC using a calibration curve (Figure S10) after 24 h incubation of amorphous PET microparticles (microPET) with PHL7, D5 or D11 (Figure 3 and Figure S11 and S12). The microPET exhibited a size of 8422 nm, polydispersity index of 0.200 and a zeta potential of-34 mV by DLS and LDA, respectively. This microPET was derived from PET nanoparticles with a size of 194 nm, polydispersity index of 0.251 and zeta potential of-38 mV by DLS and LDA, respectively. The size increase from nanoparticles to microPET occurred upon addition of the degradation buffer (1M K_2_HPO_4_ titrated with HCl to pH 8.0, 200 mM NaCl) for the enzymatic PET degradation assays.

**Figure 3.**
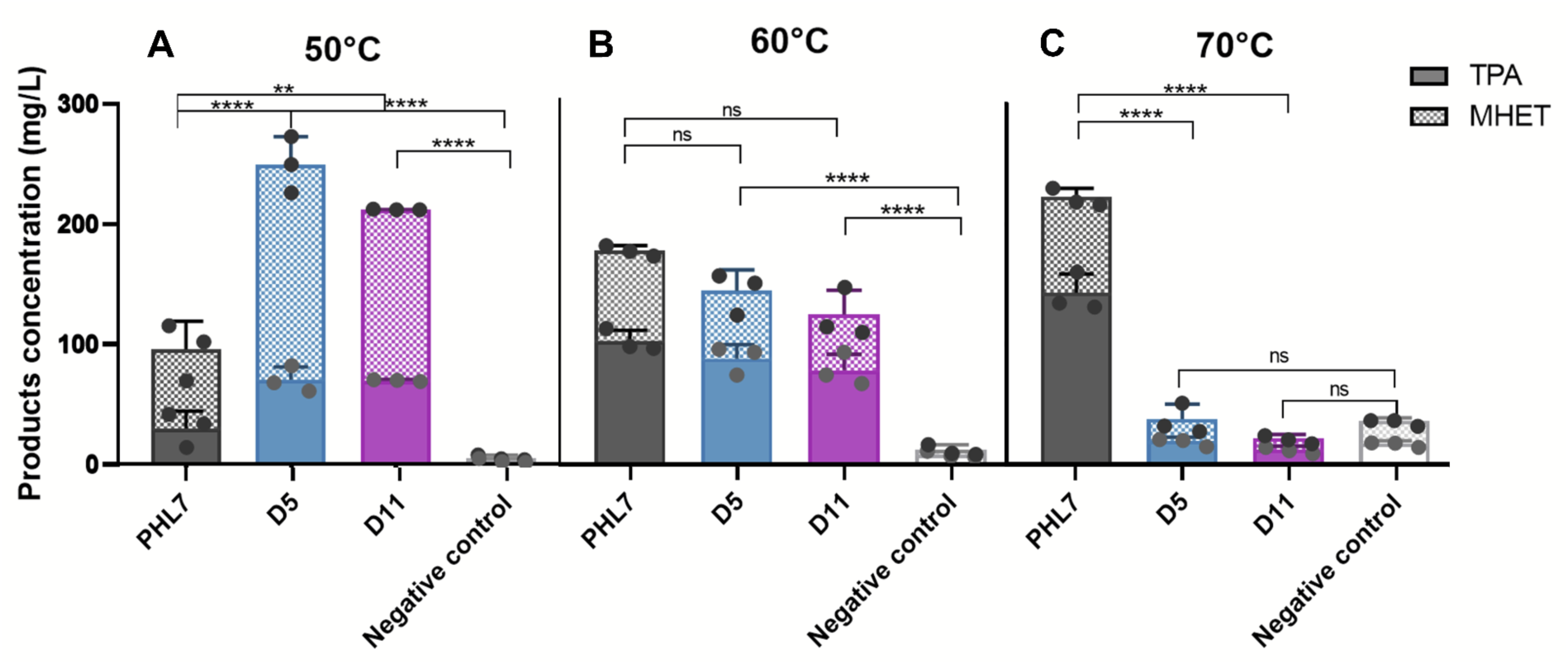
Enzymatic degradation of microPET by PHL7 and active PHL7 redesigns. Amounts of TPA (grey) and MHET (hatching) released by PHL7 and the active redesigns D5 and D11 (18.04 mg_ENZYME_/g_PET_) after a reaction of 24 h at 50°C (A), 60°C (B), and 70°C (C) in 1M K_2_HPO_4_ titrated with HCl to pH 8.0, 200 mM NaCl. The microPET degradation assays were performed in triplicates for all enzymes in all conditions. Product concentrations were estimated based on the calibration curves in Figure S10. Negative controls correspond to reactions without enzyme.

PHL7 showed higher microPET conversion rates of 43.8% after 24 h at elevated temperatures (Figure 3C and Figure S12), but conversion decreased by more than half at 50°C (Figure 3A and Figure S12). In contrast, lowering the temperature from 70°C to 50°C increased D5 and D11 conversion by 6-fold and 7-fold, respectively, going from 5.7 to 32.4% in D5 and 4.1 to 28.7% in D11 (Figure 3A, 3C and Figure S12). In fact, D5 and D11 produced total soluble products at 50°C that are comparable to PHL7 at 70°C, with a slight increase of production for D5 in comparison to the others, indicating similar PET degradation rates but at lower temperatures and with higher MHET relative to TPA (Figure S13).

At 60°C, all enzymes produced similar TPA and MHET quantities after 24 h (Figure 3B). At 50°C, D5 and D11 generated more MHET than TPA, with higher amounts of the latter compared to PHL7 (Figure 3A and Figure S12). This is relevant, because recent publications suggest that BHET and MHET may enable more efficient direct repolymerization into virgin PET. Conventional PET resynthesis requires condensing TPA and EG into BHET as the starting material for polycondensation^25^, whereas recently described MHET-initiated repolymerization strategies offer new PET resynthesis routes^26^.

To determine initial rates of PET degradation, kinetics parameters were calculated using inverted Michaelis Menten equation for all enzymes after performing reactions against PET films at 50, 60 and 70°C with varying enzyme concentrations^27^, sampled every minute for 20 min by absorbance at 240 nm (Table 3 and Figure S14). Except for ^inv^V_max_/S_0_ of D11 at 50°C, all enzymes showed increased ^inv^*V*_max_/S_0_ with temperature. PHL7 also exhibited an increase in ^inv^K_M_ with increasing temperature, whereas D5 and D11 showed a decrease in ^inv^K_M_ with temperature.

**Table 3.** Kinetic parameters of PET film hydrolysis by PHL7 and native redesigns D5 and D11 at varying temperatures.

| Temperature | 50°C |  | 60°C |  | 70°C |  |
| --- | --- | --- | --- | --- | --- | --- |
| Enzyme | $^{inv}V_{max}/S_0$<br>(10<sup>-3</sup>/min·mg) | $^{inv}K_M$<br>(μL/mg) | $^{inv}V_{max}/S_0$<br>(10<sup>-3</sup>/min·mg) | $^{inv}K_M$<br>(μL/mg) | $^{inv}V_{max}/S_0$<br>(10<sup>-3</sup>/min·mg) | $^{inv}K_M$<br>(μL/mg) |
| PHL7 | 16.14 | 1.88 | 23.19 | 27.79 | 46.67 | 35.07 |
| D5 | 8.25 | 10.88 | 15.36 | 10.12 | 22.03 | 6.29 |
| D11 | 9.11 | 14.04 | 5.36 | 11.73 | 6.92 | 7.75 |

Initial rate measurements show PHL7 exhibits higher turnover rates than D5 and D11 under saturating conditions (Table 3), mirroring observations from PCL nanoparticle assays at 50°C (Table 2). However, D5 and D11 exceed PHL7 activity in 24 h at 50°C.

These results support PHL7 inactivation at lower temperatures during the extended reaction times necessary for PET degradation. This was observed for a previously engineered PHL7 L92F/Q94Y double mutant^12^, which showed slower depolymerization at 45 and 50°C than at 60°C in 48 h experiments with cryo-ground amorphous PET films. After 24 h, PET depolymerization was 55% and 20% lower at 45°C and 50°C, respectively, compared to 60°C^14^. Initial rate measurements do not capture this inactivation, confirming a thermostability-activity tradeoff across variants.

To understand these tradeoffs, D5 and D11 sequences and structures were compared to PHL7 and analyzed using Rosetta3^28,29^. Both designs are highly conserved, with 84.56% (D5) and 84.17% (D11) sequence identity (Figure S15), corresponding to 39 and 40 substitutions, respectively, and with 20 substitutions shared between designs (Table S3). The active site is highly conserved, with the exception of two substitutions in D5 not reported in other PHL7 variants: L93T and Q95L (Figure S16). Rosetta3 analysis shows 17 D5 substitutions and 18 D11 substitutions (9 shared) exhibit more favorable energies (ΔEnergy <-1 kcal/mol) than native PHL7 residues, while 8 substitutions in each design (2 shared) show unfavorable energies (ΔEnergy > 1 kcal/mol) (Table S3). Overall, the designs exhibit slightly more favorable total energy than PHL7 (Table S4).

Analysis of D5 and D11 reveals alterations in stability-associated residues even without direct substitutions (Table S5). Both designs show systematic reduction in complex salt bridge networks (specifically those involving 3 residues) compared to PHL7 (Table S6), consistent with their experimentally measured T_m_ decreases.

To investigate the increased activity of the designs at lower optimal temperatures, molecular dynamics (MD) simulations of PHL7, D5 and D11 were performed at 50, 60 and 70°C (Figure 4 and Figure S17). Cumulative analysis of three independent 200 ns MD simulations trajectories per enzyme revealed increased active site flexibility in D5 and D11 compared to PHL7, particularly in loop regions responsible for PET binding (Figure 4). These include the W-loop, which harbors a tryptophan (W156) that clamps TPA rings via π-stacking interactions^15^, and the D-loop, which contains the catalytic aspartate (D177)^30^. Substitutions that increase loop flexibility have been shown to enhance depolymerization activity and decrease the optimal temperature in engineered PET-degrading enzymes^30,31^.

**Figure 4.**
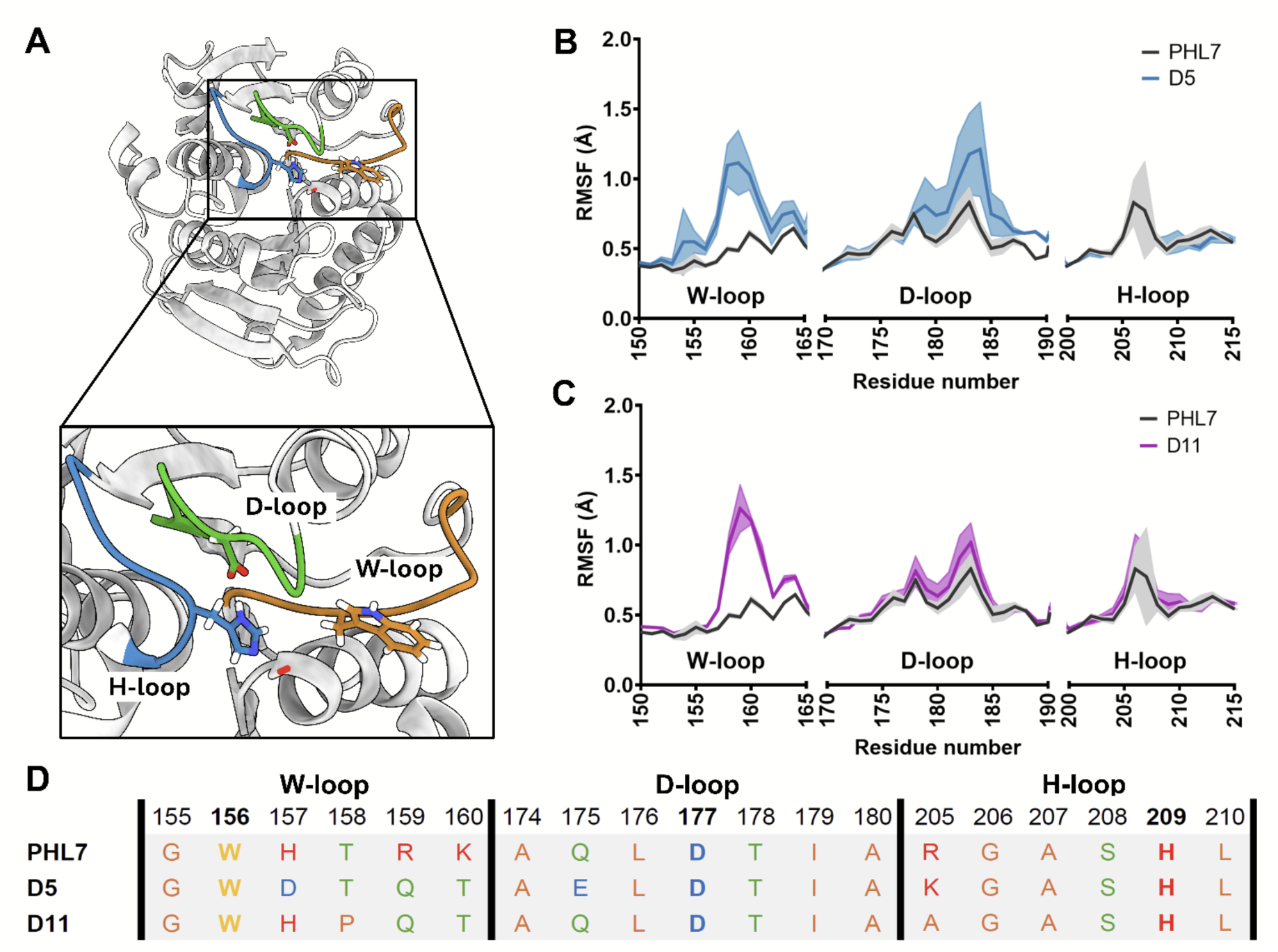
Impact of residue substitutions in the designed PHL7 variants on the active site flexibility. (A) Structure of PHL7 with the flexible active site W-loop (harboring a TPA-clamping W156 in orange), D-loop (harboring the catalytic D177 in green), and H-loop (harboring the catalytic H209 in blue). (B-C) Cα root-mean-square fluctuations (RMSF, Å) of W-loop, D-loop, and H-loop during MD simulations at 50°C for D5 (B) and D11 (C) compared to PHL7. The shaded area represents the standard deviation over 3 × 200 ns independent trajectories per simulation system. (D) Amino acid sequence of the active site loops analyzed in the MD simulations.

The MD simulations suggest that these localized destabilizations result directly from substitutions introduced near critical regions during sequence redesign. At higher temperatures, the MD simulations identified flexibility hotspots in D5 and D11 that likely promote partial denaturation, correlating with activity loss during long PET degradation reactions under these conditions (Figure S17).

D5 exhibited pronounced flexibility changes in distal regions not directly associated with the active site, particularly associated with the loop between helix α3 and strand β5 (Figure S18). This increased flexibility appears to originate not from loop mutations but from the nearby D36K substitution, which alters buried residue exposure based on Rosetta3-relaxed structures and may explain the enhanced α3-β5 loop flexibility. Similar distal flexibility effects were observed in the Antarctic PET hydrolase Mors1. Swapping its extended post-H-loop region with the equivalent LCC segment altered local flexibility in regions neighboring the active site, including the α1-β1 loop and helices η3 and α2^32^.

## DISCUSSION

This study focused on the design of more expressible, thermostable, and active variants of the PET-degrading enzyme PHL7, using ProteinMPNN and LigandMPNN in combination with structural and evolutionary information. Of 36 sequences tested, 31 were successfully purified with higher yields than PHL7, but only D5 and D11 exhibited similar PET hydrolase activity in initial high-throughput screenings. D5 and D11 showed lower thermal stability but superior depolymerization at reduced temperatures, producing similar product levels at 50°C as PHL7 at 70°C. Both generated more MHET than TPA at 50°C, recently identified as a more efficient PET repolymerization pathway^26,33^, making the strong depolymerization activity of D5 and D11 at 50°C promising for biotechnological applications^34^.

The designs present sequence changes that exhibited improved protein yields at micro-and large scales^5^. Possible explanations are related to how MPNN tools work. The model generates greater sequence-structure compatibility^17^. It tends to produce sequences with more favorable energy and lower structural frustration, which is associated with less misfolding and fewer inclusion bodies, in addition to avoiding exposed hydrophobic surfaces. In fact, ProteinMPNN described increases in yield for 73/96 designs of different proteins.

It is important to note two limitations of this study related to the potential industrial application of these enzymes. First, we did not test if the redesigned enzymes also have higher expression than the parental PHL7 in common hosts for industrial production of enzymes, such as *Bacillus* spp^35^, *Aspergillus* spp or *Trichoderma* spp^36^. However, previous works have shown that some computationally designed proteins with higher expressibility in *E. coli* also have higher expressibility in other models, including eukaryotic organisms^37^. Second, we did not test the efficiency of these enzymes in small-scale bioreactors mimicking industrial conditions and higher substrate loading, which we hope to validate in future works.

The designed enzymes exhibit an activity-stability tradeoff: reduced thermostability enabled conformational flexibility for efficient catalysis at lower temperatures, similar to mesophilic PET hydrolases^38^. Structural analysis revealed fewer salt bridge networks than native PHL7. These structural features can determine conformational dynamics and thermal stability^39^, and can distinguish psychrophilic from mesophilic enzymes^40^. MD simulations showed increased flexibility localized to PET binding and catalytic regions, associated to the mutations in each variant. An example is the Q175E mutation present in the D5 variant, which appears to confer increased flexibility in the D-loop. The D11 variant does not have this mutation, which would explain the reduced increase in flexibility compared to D5, and which appears to be associated with mutations located further away from this region. The variants present a similar behavior to *Is*PETase-inspired variants that enhanced activity at lower optimal temperatures^30,31^.

At 70°C, D5 and D11 lacked PET-degrading activity, likely due to reduced thermostability. This was unexpected, as ProteinMPNN typically generates sequences with high thermal stability. However, the inherent high thermostability of PHL7 likely complicated further improvements. Newer tools like HyperMPNN^41^ may be better suited for generating thermostable variants from already stable scaffolds. Our inability to identify thermostable designs may also stem from testing only 3% of generated sequences due to budget constraints. While this preserved activity in two designs, it precluded thorough sequence-function exploration. Directed evolution has yielded active variants with mutations at positions we conserved during our sequence redesign pipeline^42^.

This study is one of many that have sought to identify more efficient variants for PET degradation. Among the early studies of mutants, the discovery of the PHL7mut3 variant through directed mutagenesis stands out; the original paper evaluated the L93F, Q95Y, and L210T mutations individually, demonstrating an increase in stability and enzymatic activity^43^; however, when these mutations were combined, it was observed that their effects are not additive, since while Q95Y is beneficial on its own, this effect diminishes when L210T is added^44^. Other studies that employed directed evolution in conjunction with rational design^42^ succeeded in obtaining four new variants with improved properties, among which PHL7-Jemez stands out for its greater degradation capacity in bioreactors. Similarly to this work, during submission of our manuscript a study was published with a new four-step pipeline that combines computational design based on PROSS with rational mutagenesis, generating variants with higher melting temperatures and improved activity at lower phosphate concentrations (0.1 M)^44^. This is an important characteristic of PHL7 variants, since the need for a high phosphate concentration is related to enzyme stability, particularly in maintaining salt bridges within the structure. The variants obtained in this study feature new, internal salt bridges, making them less dependent on external salts; they also exhibit a reduction in surface negative charge, thereby decreasing the repulsion between acidic residues when ionic strength is low. This is not observed in the variants presented in our study, which makes them still dependent on high salt concentrations.

In the future, it would be interesting to consider new sequence selections. When clustering the sequences, diversity was prioritized over functionality. There are new high-throughput computational protocols that use MD simulations to filter out structures by binding to the PET molecule^45^. This would allow us to select sequences with greater confidence in potential functional enzymes. Furthermore, given the nature of the study and considering the subsequent results, it is likely that our activity screening was limited. It is possible that other sequences exhibited activity at either lower or higher temperatures, but due to the limited testing of sequences at temperatures near 70°C, sequences that could potentially degrade at other temperatures than those presented in this study were missed.

In summary, this work demonstrated that deep learning-guided design and activity screening are effective in generating PET-degrading enzymes with improved protein titers from heterologous expression and high activity at moderate temperatures, providing practical guidance for navigating sequence selection strategies when computational throughput exceeds experimental capacity. Thus, these designed enzymes may constitute strong candidates for usage in industrial setups due to their high efficiency at mild temperatures, which would also reduce evaporation losses. Also, their high PET depolymerization efficiency at 50°C makes them more compatible with other processes, such as whole-cell biocatalysis using moderately thermophilic hosts^46,47^. These results indicate that enhanced low-temperature activity can emerge at the expense of thermostability, revealing a designable activity–stability balance. All of this brings us closer to improvements in different types of biorecycling systems, which are essential to enable scalable and economically viable circular PET economies.

## METHODS

### Sequence redesign of PHL7

Rosetta3^28^ relaxed^23^ chain A of the crystal structure of PHL7 bound to TPA (PDB ID 8BRB) was selected as the input backbone for ProteinMPNN^17^ and LigandMPNN^18^. To maintain the catalytic activity of the designs, the active site residues were chosen based on their location at a 6 Å radius from TPA in the structure and fixed during the sequence redesign process.

This information was combined with fixed residues based on their sequence conservation, an evolutionary information that was generated using 2 multiple sequence alignments (MSAs) of PETases as inputs: (i) PET PAZy, which includes sequences from the Plastics-Active Enzymes Database^48^, and (ii) PET selected, 9 sequences of PETases with known activity tested by our lab. The MSAs were provided to hhblits^21^ to generate four iterative searches against Uniref30^20^ (accessed January 8th, 2024) at e-value cutoffs of 1e-50, 1e-30, 1e-10, and 1e-4. The same cutoffs were applied to filter the generated MSAs based on the resulting e-values after realignment using a Maximum ACcuracy (MAC) algorithm by hhblits, which were used to generate a new MSA. With this, a total of 16 MSAs were generated as a result.

Amino acids were then filtered and selected in all MSAs based on their frequency at each position, to rank them at 30%, 50%, and 70% of sequence conservation. With this, we obtained 48 different residue position groups from all 16 MSAs at 30, 50, and 70% sequence conservation to be fixed during the sequence redesign process, plus an additional set corresponding to only fixing the active residues (Table S1).

All these fixed residue position groups were employed as input for 98 different fixed backbone sequence redesign campaigns using ProteinMPNN^17^ and LigandMPNN^18^. In these campaigns, the active site and residues at different sequence conservation cutoffs from the 49 aforementioned groups were fixed, and cysteines were excluded from the sequence redesign process, except for those already present on PHL7 and forming disulfide bonds. For both ProteinMPNN and LigandMPNN, a conservative temperature of 0.05 and a Gaussian noise of 0.10 were used. A total of 12 sequences were built for each group of conserved positions, generating a total of 1176 sequences (588 with ProteinMPNN and 588 with LigandMPNN). Afterwards, repeated sequences were eliminated.

Structures for each redesigned sequence were predicted by ESMFold^22^ using 12 recycling steps. The structures were filtered using a Cα RMSD <1.5 Å against the crystal structure of PHL7 and a predicted local distance difference test (pLDDT, a metric of the overall confidence of the predicted protein structure)^49^ >85. The structures were relaxed using Rosetta3^28^ and the FastRelax protocol^23^ and scored based on their total energy (in kcal/mol). Designs with a total energy >-200 kcal/mol were eliminated.

Despite all these filters, less than 100 sequences were eliminated through this pipeline. Therefore, to reduce the number of sequences to test, a Levenshtein matrix was used to cluster the sequences according to their similarity and select 36 representative sequences from each enzyme cluster for experimental testing. The initial methionine missing from the sequence was added to the selected sequences.

### Cloning and expression of PHL7 and sequence redesigns

Codon-optimized eBlocks gene fragments encoding PHL7 and the 36 PHL7 sequence redesigns were purchased from Integrated DNA Technologies, USA. A Golden Gate cloning protocol was performed using the BsaI-HFv2 enzyme (New England Biolabs, USA) to insert the fragments into a custom pET29b vector designed for Golden Gate cloning (pAIP4), which adds a 10X Histidine tag and a SUMO tag at the N-terminal of the enzyme of interest to then express them in the cytoplasm (Addgene Plasmid #233686). Cloned designs were transformed by heat shock into chemically-competent *E. coli* BL21(DE3) cells containing a custom helper plasmid overexpressing human *PDIA1* (a protein disulfide isomerase) and yeast *ERV1* (a sulfhydryl oxidase) to facilitate correct disulfide formation in the *E. coli* cytosol, equivalent to the CyDisCo plasmid^50^, and recovered on agar plates.

To check if the correct fragment was cloned, expression tests were performed for each design. Colonies were selected and inoculated into 200 μL autoinduction media (ZYM-5052 media with 100 μg/mL kanamycin, 34 μg/mL chloramphenicol, and 0.01% v/v Antifoam Y-30 Emulsion) to grow overnight. Sodium dodecyl sulfate-polyacrylamide gel electrophoresis (SDS-PAGE) was performed the next day. Enzymes exhibiting the expected molecular weight were selected and expressed into 2 mL of autoinduction media in a 24-well deep plate at 25°C overnight. The next day, cells were precipitated and frozen until used. Cells of the successfully cloned enzymes were stored as glycerol stocks for further work.

### High-throughput enzyme purification

Cells were thawed completely at room temperature to be resuspended with 2 mL of buffer A (50 mM Na-phosphate buffer pH 8, 200 mM NaCl) supplemented with 0.25 mg/mL lysozyme, 25 U/mL Turbonuclease, and 1x EDTA-free Protease Inhibitors. Lysates were sonicated for 5 min on ice following a protocol of 30s ON, 30s OFF, 55% amplitude. Supernants were collected by centrifugation at 3900 rpm for 10 min and incubated with Ni-NTA magnetic beads, already equilibrated with buffer A, for 30 min at 350 rpm. The magnetic beads were washed 4 times using 1 mL of buffer A. To elute the proteins, 600 μL of elution buffer (buffer A supplemented with 600 nM *Cth* SUMO protease^51^) was added to each plate overnight at 350 rpm at room temperature.

The protein yield of all purified enzymes was measured based on fluorescence using the Qubit™ Protein and Protein Broad Range (BR) Assay Kits (Thermo Fisher Scientific, USA). To determine the purity of the samples, analytical size exclusion chromatography (SEC) was performed using an AdvanceBio SEC HPLC column (300Å 4.6 x 300 mm, 2.7 μm; Agilent Technologies, USA) on a Nexera HPLC System Controller SCL-40 (Shimadzu Scientific Instruments, USA). Samples were supplemented at a rate of 0.35 mL/min using buffer A as the mobile phase.

### Enzyme purification on a large scale

PHL7 and active sequence redesigns were taken from the glycerol stocks to grow them in 2 mL of LB medium overnight at 37°C. This overgrown medium was used to inoculate 1 L of 1% 2xYT medium with its respective antibiotics (100 μg/mL kanamycin, 34 μg/mL chloramphenicol). Once an optical density at 600 nm (OD_600_) of ∼0.6 was reached, the culture was cooled and induced overnight with 1 mM isopropyl β-D-1-thiogalactopyranoside (IPTG) at 21°C and 150 rpm. Cells were sedimented by centrifugation at 5000 rpm for 30 min and then frozen until used.

The cell pellet was thawed and resuspended with 20 mL of buffer A supplemented with 0.25 mg/mL lysozyme and incubated for 20 min at room temperature. The sample was lysed by sonication (4s ON, 4s OFF, 40% amplitude) and then centrifuged for 30 min at 12000 rpm. The clarified lysate was recovered and supplemented with Ni-Sepharose resin (Ni Sepharose 6 Fast Flow, Cytiva, USA), previously equilibrated with buffer A, and incubated for 1 h at room temperature. The resin was washed 3 times with buffer B (50 mM Na-phosphate buffer pH 8, 200 mM NaCl, 20 mM imidazole) and a final time with buffer A. To elute the proteins, 2 mL of elution buffer (buffer A supplemented with 600 nM *Cth* SUMO protease) was added to the resin and incubated overnight at room temperature. The protein was eluted from the resin by washing 4 times with 2 mL of buffer A. Lastly, the protein concentration was determined by Bradford assay^52^, and its purity was confirmed by SDS-PAGE.

### Determination of secondary structure of high-throughput purified enzymes

To determine the secondary structure and correct folding of the variants, Circular Dichroism (CD) was performed. CD spectra were measured with a Chirascan Q100 CD spectrometer (Applied Photophysics Limited, UK) equipped with a Peltier thermostatic system under constant nitrogen flux with a 0.1 cm quartz cuvette. The proteins were normalized to get 0.2 mg/mL and measured at 25°C between 190 and 250 nm.

### Thermal stability measurements of high-throughput purified enzymes

To determine the thermal stability of PHL7 and the successfully purified PHL7 sequence redesigns, differential scanning fluorometry (DSF) measurements were carried out using the GloMelt™ Thermal Shift Protein Stability Kit (Biotium, USA) on the QuantStudio3 instrument (Applied Biosystems, USA). Instructions were followed according to the kit, and solutions to generate a 20 μL final sample volume were prepared. This includes: 2 μL of 200X GloMelt dye diluted at 10X on PBS (final concentration 1X), 2.5 μL of ROX reference dye (40 μM) at 400 nM (final concentration 50 nM), 10 μL of protein normalized at 1 μg/μL (0.5 μg/μL), and 5.5 μL of PBS buffer. The final volume was added in a qPCR plate with an optical seal, and the following program was run: 25°C for 5 min, followed by increasing the temperature to 99°C at a rate of 0.03°C/s, and then decreasing the temperature to 25°C. The T_m_ was calculated as the lower peak for the negative derivative of the fluorescence (-dRFU/dT). Experiments were performed with only one replicate per enzyme.

### Fast screening of PET degradation activity using high-throughput purified enzymes

When PET is enzymatically depolymerized, small soluble compounds such as BHET, MHET and TPA are produced. The presence of these products containing free carboxyl groups can produce a drop in pH in a solution with low buffering conditions^34^. An easy way to measure pH decrease is using halochromic compounds such as phenol red, which presents two absorbance peaks at 550 and 430 nm for red and yellow colors at high and low pH, respectively^34^. In addition, the soluble products generate an increase in absorbance at 240 nm due to the presence of the same number of carbonyl groups^53^.

A fast and high-throughput PET degradation screening test was thus designed to measure both the absorbance at 240 nm due to the presence of soluble products and the pH change they produce in the reaction solution in the presence of phenol red. For this assay, reactions were performed in 96-well PCR plates containing 10 µL of enzyme from normalized stock solutions at 90.5 μg/mL (final concentration of 9.05 μg/mL), 6.9 µL of a solution of 2.17 mM phenol red on PBS (final concentration 0.15 mM), 10 µL of buffer containing 1 M K_2_HPO_4_ titrated with HCl to pH 8.0 and 200 mM NaCl (final concentration 100 mM K_2_HPO_4_, 20 mM NaCl, pH 8.0) and 73.1 µL of milliQ water. A small post-consumer amorphous PET film (half circles of 6 mm of diameter) was added to the plate. Plates were sealed and incubated at 60, 70 and 80°C for 16 h. Measurements of absorbance at 240 nm (BHET, MHET and TPA absorbance), 430 nm (yellow absorbance due to acidification), and 550 nm (red absorbance due to basicity) were taken before and after the experiment.

### Thermal stability of PET hydrolases produced at large scale using DSF

To determine the thermal stability of PHL7 and the active PHL7 sequence redesigns after purification at large scale, DSF measurements were carried out using SYPRO Orange 5000X (Thermo Fisher Scientific, USA) on the QuantStudio3 instrument. Samples were prepared on a 20 μL final volume that includes: 19 μL of a 0.5 mg/mL enzyme solution in buffer A and 1 μL of a previously diluted solution of 5000X SYPRO Orange at 50X in DMSO. The volume was added in a qPCR tube with an optical seal, and the following program was run: 5°C for 2 min, followed by increasing the temperature to 99°C at a rate of 0.01°C/s, and then decreasing the temperature to 25°C. The T_m_ was calculated as the lower peak for the negative derivative of the fluorescence (-dRFU/dT). DSF was performed in triplicates.

### Determination of optimal reaction temperature using polycaprolactone nanoparticle degradation assays

Polycaprolactone (PCL) nanoparticles were prepared by nanoprecipitation and solvent evaporation, similarly to previous works, but with minor modifications described below^54^. Briefly, 30 mL of acetone was supplemented with 250 mg of PCL (average molecular weight ∼14000 g/mol, Sigma-Aldrich, USA) and heated to 50°C. The solution was added dropwise into 100 mL of MilliQ water previously heated and constantly stirred. The final solution was kept under agitation at a temperature of 50°C until the acetone had completely evaporated. The physicochemical characterization of the PCL nanoparticles was performed using a Malvern Zetasizer dynamic light scattering (DLS) and laser doppler anemometry (LDA) instrument (Malvern Instruments, UK).

The hydrolysis rates of PCL nanoparticles by PHL7 and the active PHL7 sequence redesigns were determined by the change in turbidity at OD_600_ as an indirect measure of particle depolymerization. To determine the optimal degradation temperature, reactions were measured at a final time of 10 min. The reactions contained a mixture of 50 µL of PCL nanoparticle suspension (stock concentration of 2.6 mg/mL), 25 µL of degradation buffer (1M K_2_HPO_4_ titrated with HCl to pH 8.0, 200 mM NaCl), and a final enzyme concentration of 25 µg/mL from a protein solution in buffer A, supplemented with buffer A up to a final volume of 100 µL (19.2 mg_ENZYME_/g_PCL_). This composition ensured an initial turbidity at OD_600_ of ∼0.8-1.0 for the PCL nanoparticle degradation assays. Temperatures between 25°C and 70°C were tested in 5°C increments. All measurements were made in triplicate in an EPOCH2 spectrophotometer (BioTek Instruments, USA). The final degradation of PCL nanoparticles after 10 min of incubation was calculated using the following formula:

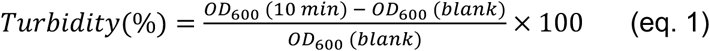

### Kinetic parameters of PCL hydrolysis

To determine the kinetic parameters of PCL nanoparticle hydrolysis by PHL7 and the active PHL7 sequence redesigns, assays were performed at 50°C using a reaction mixture in a final volume of 100 µL, containing 60 µL of PCL nanoparticle suspension (stock concentration of 2.6 mg/mL), 25 µL of degradation buffer (1M K_2_HPO_4_ pH 8.0, 200 mM NaCl), and enzyme in varying concentrations between 1 and 30 µg/mL in buffer A. Again, this mixture was made in order to achieve an initial turbidity at OD_600_ of ∼0.8-1.0 for the PCL nanoparticle degradation assays. The tests were performed in 96-well microplates, measuring the decrease in turbidity at OD600 up to 30 min in an EPOCH2 spectrophotometer (BioTek Instruments, USA). The tests were performed in triplicate. Only the initial velocity was taken for the calculations, determined by the linear region of the graphs in the decrease in turbidity using the kinetic parameters determined as a pseudo-first-order kinetic equation^24^:

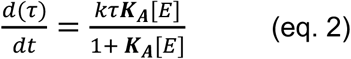

With *τ* as the turbidity of a nanoparticle suspension, *t* as the reaction time, *k_τ_* as the hydrolysis rate constant based on the turbidity change, K_A_ as the adsorption equilibrium constant, and [E] as the enzyme concentration. Statistical significance was determined using a Two-way ANOVA test.

### Determination of optimal reaction temperature for PET degradation

The hydrolysis rates of PET by PHL7 and the active PHL7 sequence redesigns were determined by the increase in soluble products after incubation of amorphous PET films with the enzyme. To determine the optimal temperature for PET depolymerization, measurements were taken at 30 min at temperatures ranging 25-70°C using a mixture of 2.5 mm x 2.5 mm amorphous PET films (250 μm thickness; product number ES301445, Goodfellow, Germany), 25 µL of degradation buffer and 30 µg/mL of enzyme solution in buffer A. The degradation products were measured by absorbance at 240 nm in a EPOCH2 spectrophotometer (BioTek Instruments, USA). Assays were performed in triplicates.

### Preparation of PET nanoparticles

PET nanoparticles were prepared following previous publications^55^, with modifications regarding the source material, using in this case granular PET (Sigma-Aldrich, USA). To produce PET nanoparticles, 1 g of granular PET was dissolved in 10 mL of trifluoroacetic acid (TFA) 90% v/v, which was left under stirring at 800 rpm and 50°C until complete dissolution, after which the mixture remained under room temperature overnight without stirring. The next day, the dissolution was kept under stirring at 1000 rpm and 30°C for 30 min, upon which 10 mL of TFA 20% v/v was added dropwise, and the mixture was maintained under stirring for 2 h and then left undisturbed overnight at room temperature. Finally, the dissolution was centrifugated for 1 h at 2500 g, and the supernatant was discarded, the pellet was resuspended in 100 mL of sodium dodecyl sulfate (SDS) 0.5% v/v and sonicated with an amplitude of 25% for 1 min with 10 s ON/OFF pauses to aid particle dispersion. Subsequently, the PET nanoparticles were centrifugated for 1 h at 5000 rpm and 35°C, after which the supernatant was discarded, the pellet was resuspended in milliQ water and continuously sonicated for 10 min. The physicochemical characterization of the PET nanoparticles was performed using a Malvern Zetasizer DLS and LDA instrument (Malvern Instruments, UK).

### Enzymatic hydrolysis of microPET

To analyze the hydrolysis of PET microparticles (microPET) by PHL7 and the active PHL7 sequence redesigns, we measured the increase in BHET, MHET and TPA during the incubation of these particles with the enzymes via high-performance liquid chromatography (HPLC). For the reactions, a mixture containing 50 µL of PET nanoparticle solution (stock concentration of 8.1 mg/mL) and 18.04 mg_ENZYME_/g_PET_ was completed up to a final volume of 500 µL with degradation buffer (0.41 mg PET). This degradation buffer led to the formation of microPET starting from the PET nanoparticles generated as described before, which were thus characterized by DLS using a Malvern Zetasizer DLS instrument (Malvern Instruments, UK). The mixture was incubated at 50, 60, and 70°C for up to 24 h with constant agitation (700 rpm). The enzymatic assays were performed in triplicates.

For the quantification of the PET degradation products MHET, BHET, and TPA, these were obtained from the supernatant after centrifuging and filtering the samples using a 0.22 µm syringe filter. Their concentration was determined using a Jasco LC-4000 HPLC with autosampler (Jasco Inc., Japan) equipped with a C18 column (Eurosper II 100-5; 150 × 2 mm with pre-column, Knauer Wissenschaftliche Geräte GmbH, Germany) at a flow rate of 1.2 ml/min. The mobile phase consisted of 19:81% acetonitrile:formic acid (0.1% v/v). The sample was diluted 20 times and 10 µL of the dilution were injected, detecting the separation of all products at a wavelength of 260 nm. Commercially available TPA (Sigma-Aldrich, USA), BHET (Sigma-Aldrich, USA), and MHET (Ambeed, USA) were used as standards. Statistical significance was determined using One-way and Two-way ANOVA tests.

### Kinetic parameters of enzymatic PET hydrolysis

To determine the kinetic parameters of enzymatic PET hydrolysis, the soluble degradation products for PHL7 and the active PHL7 sequence redesigns were collected every 1 min for up to 20 min during PET degradation reactions at 50, 60, and 70°C. To do this, the reactions were carried out in a final volume of 50 µL containing 2.5 mm x 2.5 mm PET film (Goodfellow, Germany), 25 µL of degradation buffer, and enzymes in varying concentrations from 1 to 40 µg/mL. Tests were performed in 96-well PCR plates in triplicates, using one plate for each time point. The enzymatic reaction was stopped by placing the plate in ethanol at-80°C. The degradation products were measured for each time point at 240 nm using the EPOCH 2 spectrophotometer (BioTek Instruments, USA). The initial velocity was calculated considering the linear region of this data, which were used to obtain the kinetic parameters via the Inverse Michaelis Menten equation^27^:

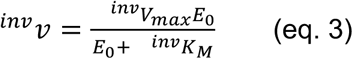

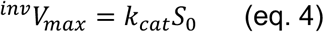

where ^inv^*v* is the initial velocity, E_0_ are gradually increasing enzyme concentrations, ^inv^K_M_ is the enzyme concentration at the half-saturation point, ^inv^*V*_max_ is the maximal rate at substrate saturation, *k*_cat_ is the catalytic turnover and S0 is the mass load of a substrate.

### Calculations of energetic and physicochemical parameters with Rosetta3

To calculate energetic and physicochemical parameters of PHL7 and the PHL7 sequence redesigns of interest, Rosetta3^28^ was used in its PyRosetta version^29^. The structures were relaxed using the FastRelax protocol^23^ and then the total Rosetta score and per-residue values in kcal/mol were calculated using a custom-made python script^56^. To obtain the number of salt bridges in each enzyme, the *debug_rosetta_salt_bridges* module was used. Python scripts were used to obtain detailed information on the position of these bonds, including a sum of criteria that combine distances and probable amino acids.

### Molecular Dynamics Simulations

Molecular Dynamics (MD) simulations were performed for PHL7 and its sequence redesigns using the AMBER 2024 simulation package^57^ with the ff19SB force field^58^ and the OPC water model^59^. Protein structures were protonated at pH 8.0 using the PlayMolecule server^60^, with protonation states manually verified for each enzyme. Each system was solvated in a truncated octahedral water box with a 1.5 nm padding distance and neutralized with Na⁺ or Cl⁻ counterions. A protocol of 2 energy minimization steps was used. The first minimization step used a steepest descent method with position restraints on water and ions, and the second one without any restraints. This was followed by 3 equilibration steps, the first one corresponding to 150 ps of heating to the target temperatures of 323, 333, and 343 K (50, 60, and 70°C) with positions restraints on the protein, the second to 1 ns of temperature and pressure equilibration at 1 bar while restraining only the protein backbone atoms, and a final 10 ns equilibration step without restraints at constant temperature and pressure.

Finally, NPT production runs of 200 ns at 323, 333, and 343 K (50, 60, and 70°C) and 1 bar were run, with 3 independent simulations per condition. Temperature and pressure were maintained using a Langevin thermostat (collision frequency = 1 ps⁻¹)^61^ and a Berendsen barostat (pressure relaxation time = 1 ps)^62^. The simulations employed a 2 fs time step, with bonds involving hydrogen constrained via the SHAKE algorithm^63^, a 10 Å cutoff for non-bonded interactions and particle mesh Ewald method^64^ for long range electrostatics. Production MD runs were analyzed based on the Cα root mean square fluctuations (RMSF) to determine potential changes in the local dynamics of different regions of these enzymes.

### Boltz-2 co-folding structure predictions

Predictions of D5 and D11 with TPA were performed using Boltz-2^65^, a deep-learning–based protein structure prediction framework that supports protein-ligand predictions. The amino acid sequences of D5, D11, and the SMILES of TPA were provided as input for Boltz-2 predictions using default settings: 5 independent model initializations, 1 diffusion sample, 200 sampling steps and 3 recycling steps, and using the MMseqs2 server^66^ to retrieve a MSA for the protein-ligand structure predictions. The best ranked model was used for analysis of active site residues surrounding the TPA moiety.

## SUPPORTING INFORMATION

SDS-PAGE and high-throughput post-consumer amorphous PET degradation screening assays of purified PHL7 and sequence redesigned enzymes at micro-scale; CD curves at 25°C and DSF curves to determine thermal stability of the purified enzymes at micro-scale; large scale protein purification results and DSF thermal stability assays, determination of optimal temperatures and initial rates of degradation against PCL nanoparticles; 24 h degradation assays and initial rates of degradation against microPET and PET films, including HPLC calibration curves for quantification of TPA, MHET and BHET; computational sequence variability and local structure flexibility analysis; groups of residues fixed in PHL7 during sequence redesign using ProteinMPNN and LigandMPNN, amino acid sequences of the 36 selected redesigned enzymes for experimental characterization; and energetic and structural analysis of PHL7 and active designs D5 and D11 using Rosetta3 (PDF).

Raw data of the CD analysis at 25°C, DSF assays from proteins purified at micro-scale and 1L scale, HPLC calibration curves using TPA, MHET and BHET, HPLC product quantification after 24 h degradation of microPET using PHL7 and redesigns D5 and D11, Rosetta3 relaxed structures of PHL7 used as input structure for sequence redesign and of designs D5 and D11, ESMFold structures of designs D5 and D11, and Boltz-2 predictions of designs D5 and D11 bound to TPA can be found on Zenodo, and are accessible at https://doi.org/10.5281/zenodo.18777210.

## ACKNOWLEDGMENTS

This research was funded by the International Centre for Genetic Engineering and Biotechnology (ICGEB) through its Collaborative Research Program (CRP/CHL23-02), the ANID Millennium Science Initiative Program (ICN17_022), the Homeworld Collective Garden Grants and the AVANZA UC project from Pontificia Universidad Católica de Chile (AV25297). AG, VE and JA were supported by ANID doctoral scholarships (PFCHA 21220450, 21231144 and 21231296). Powered@NLHPC: This research was partially supported by the supercomputing infrastructure of the NLHPC (CCSS210001). F.C.Z. and F.D-O. would like to thank Dr. José Vicente González-Aramundiz (Facultad de Química y de Farmacia, Pontificia Universidad Católica de Chile) for providing access to the Zetasizer Nano ZS (Malvern Instruments, Malvern, UK).

## AUTHOR CONTRIBUTION

A.G., V.E, F.D-O., J.A., F.C.Z., E.N.C.N., B.A.M. and C.A.R-S. conceived and planned the experiments. A.G., V.E, F.D-O. and J.A., carried out the experiments. A.G. planned and carried out the simulations. A.G. and C.A.R-S wrote the manuscript with support from F.D-O., F.C.Z., C.D.B and B.A.M. C.A.R-S, C.D.B and B.A.M. supervised the project. All authors read and approved the final manuscript.

## SUPPORTING INFORMATION

Supporting File S1. Includes Figures S1–S18 and Tables S1-S6.

