## Supporting Information for "MPNN-guided redesign of PET hydrolases with enhanced catalytic activity below the PET glass transition temperature"

This Supplementary Information contains:

- 18 Supplementary Figures
- 6 Supplementary Tables

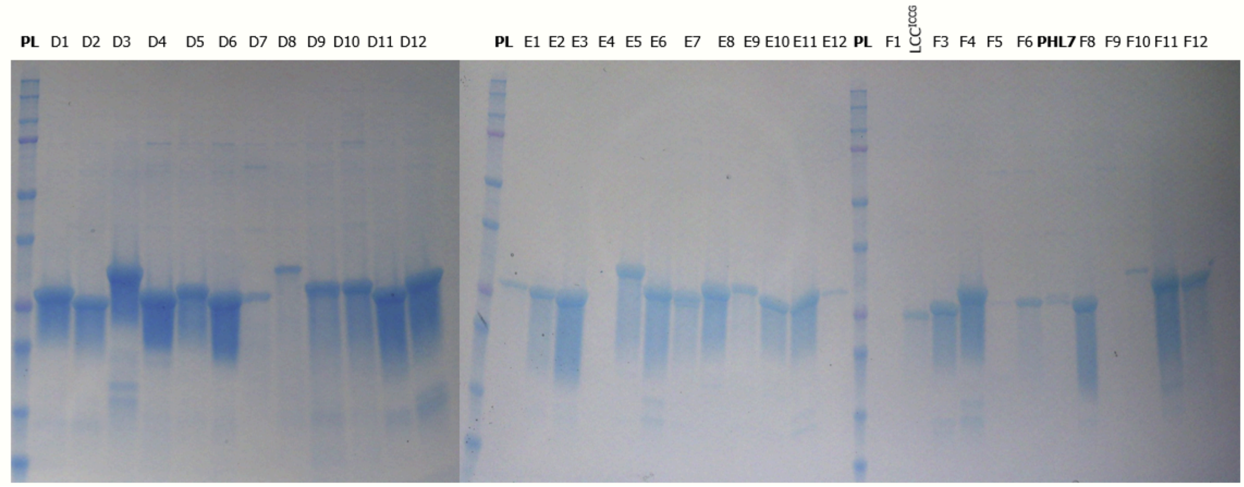


**Figure S1. Protein purification of all 36 PHL7 sequence redesigns.** SDS-PAGE gels after high-throughput purification using Ni-NTA magnetic beads. Designs E4, F1, and F9 were not observed after high-throughput purification, either due to negligible protein overexpression in the soluble fraction or low affinity to the Ni-NTA magnetic beads. F2 and F7 were unable to express. The recombinant expressions of LCC-ICCG and PHL7 were utilized as positive controls. The labels on top of each SDS-PAGE correspond to the names of the designed enzymes in each lane. PL: protein molecular weight ladder.

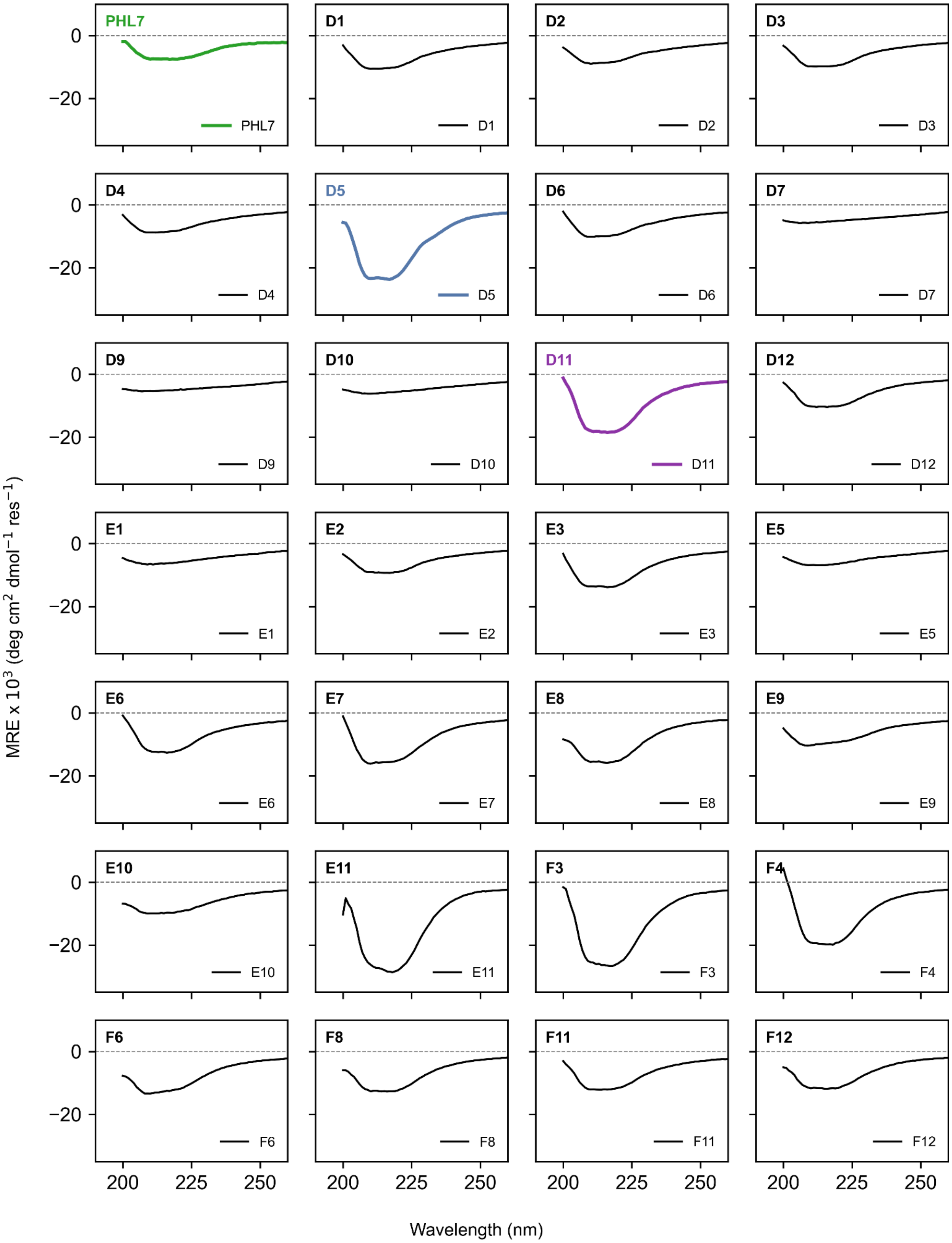


**Figure S2. Secondary structure measurements of the successfully expressed sequence redesigns of PHL7.** Circular dichroism (CD) was measured between 190 and 250 nm at 25°C, including PHL7 (green). Active designs D5 and D11 are shown blue and purple.

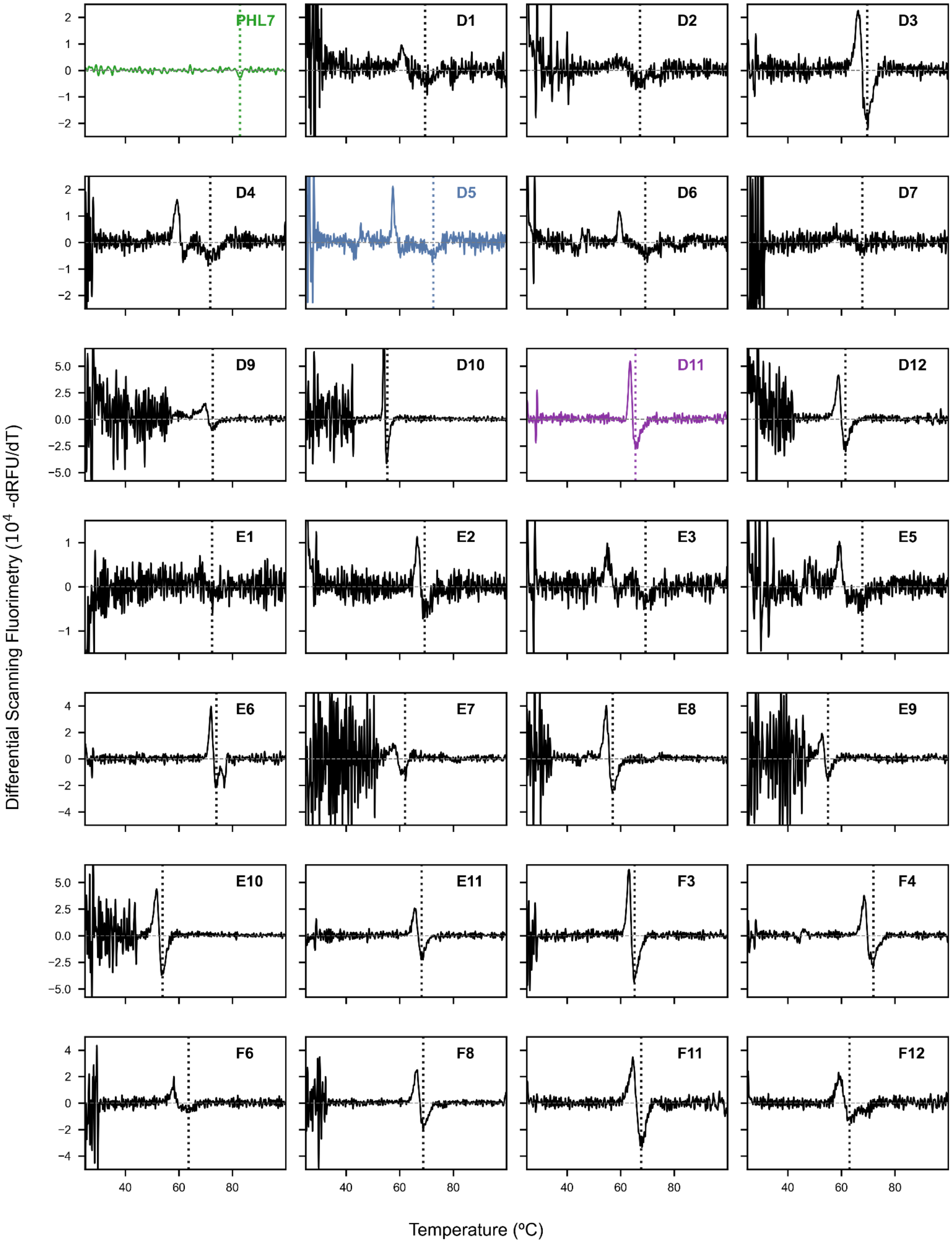


**Figure S3. Determination of the $T_m$ of redesigns by DSF.** First derivative of the ratio of GloMelt™ increasing fluorescence at 494 nm and 518 nm upon increasing temperature from 25°C to 100°C. Active designs are shown in blue and purple. DSF experiments were performed in one replicate.

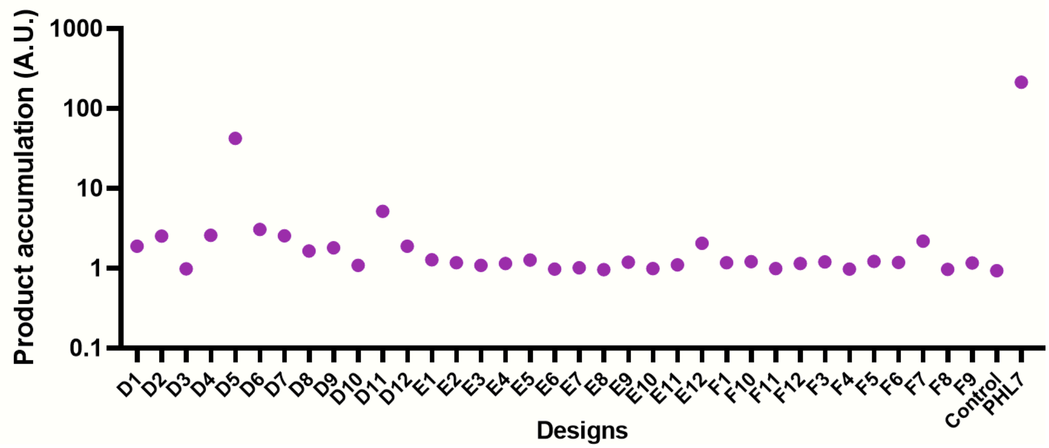


**Figure S4. Screening for PET degradation using post-consumer PET.** Degradation products were measured at 240 nm after 16 hrs of incubation with small pieces of post-consumer amorphous PET at 70°C. Quantification of product accumulation is shown in logarithmic scale. The control corresponds to a reaction without adding enzyme. Only one replicate was performed per enzyme.

.

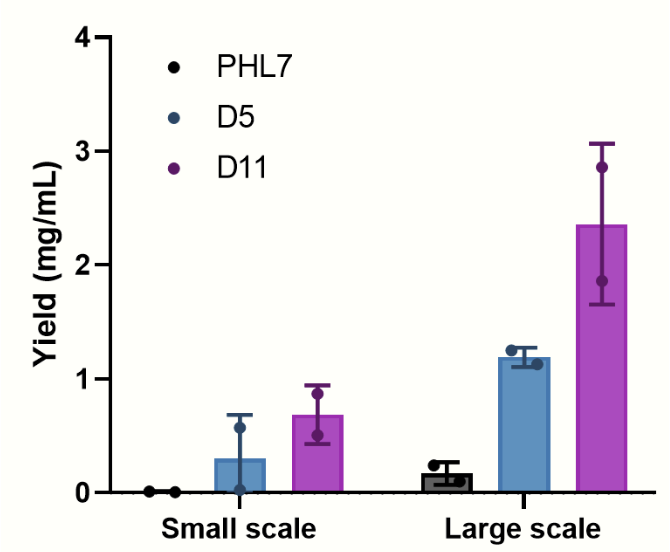


**Figure S5. Increase in protein yield between PHL7 and the native redesigns D5 and D11 at small and large scales.** Recombinant overexpression and purification of PHL7 and native redesigns D5 and D11 were performed in 2 mL (micro-scale) and 1 L (large scale). Enzyme purifications were performed in triplicate.

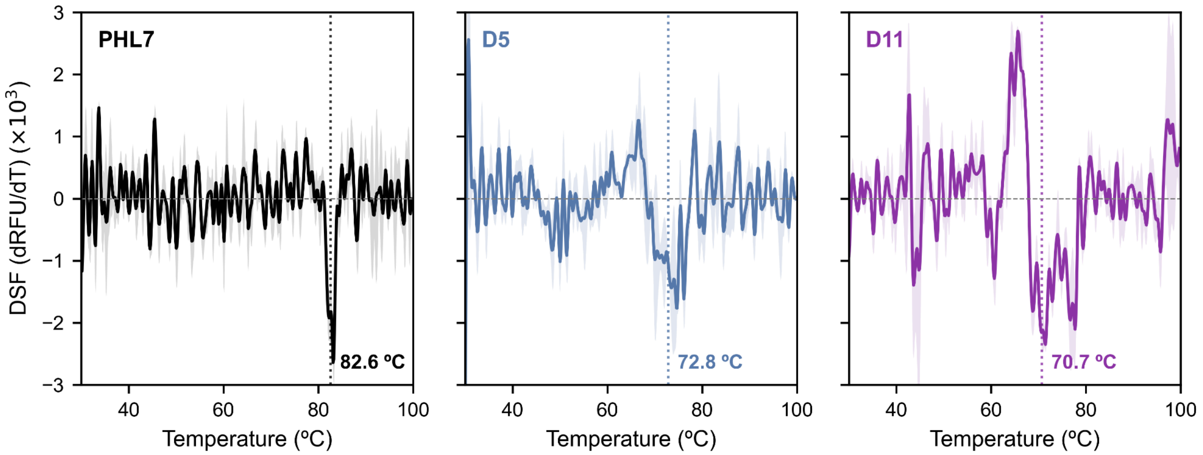


**Figure S6. Determination of the Tm of PHL7 and native redesigns D5 and D11 by DSF.** First derivative of the ratio of SYPRO orange increasing fluorescence at 494 nm and 518 nm upon increasing temperature between 25°C and 100°C for PHL7 (A) and active designs D5 (B), and D11(C). DSF experiments were performed in triplicates.

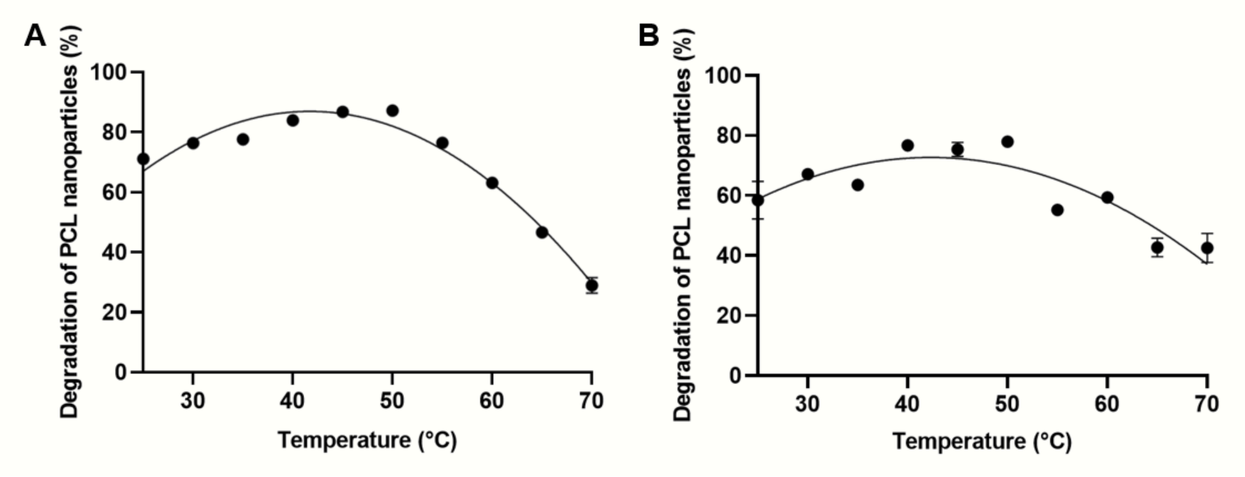


**Figure S7. Optimal temperature curves of PCL nanoparticle degradation by active PHL7 native redesigns.** Relative hydrolysis rates of PCL nanoparticles by D5 (A) and D11 (B) at temperatures ranging from 25º to 70ºC. Experiments were performed in triplicate.

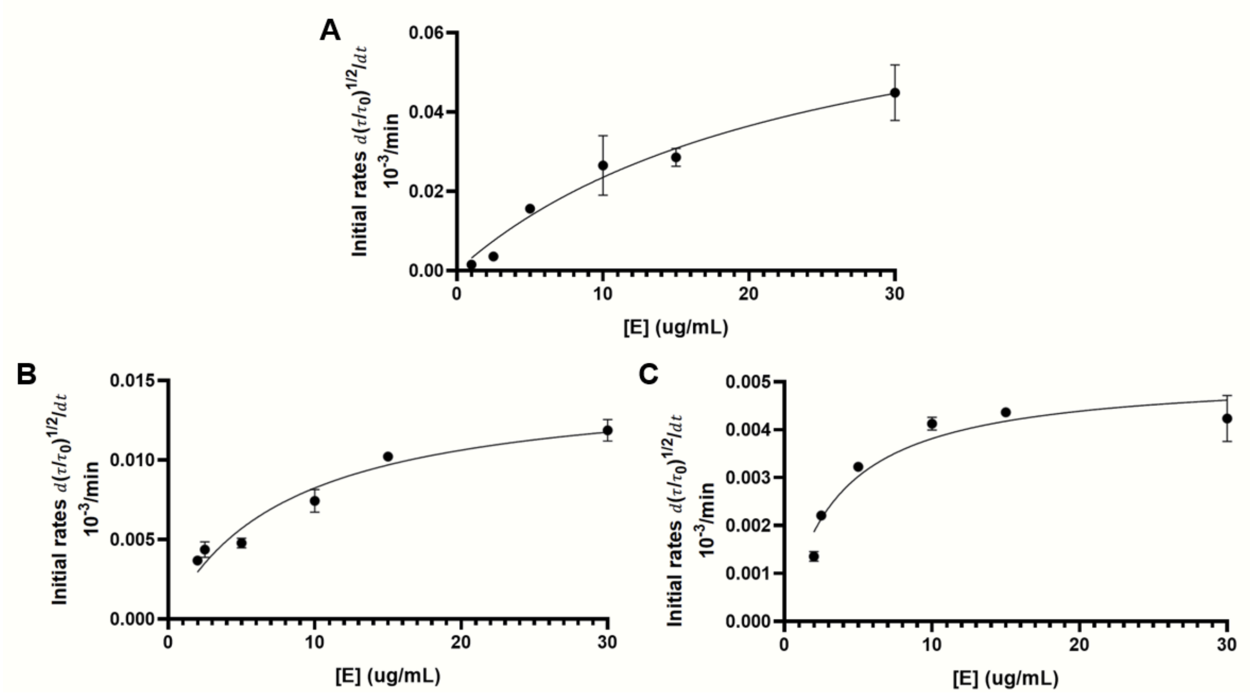


**Figure S8. Initial PCL nanoparticle hydrolysis rates by PHL7 and active redesigns D5 and D11 at varying enzyme concentrations.** Enzymatic PCL nanoparticle hydrolysis by PHL7 (A), D5 (B), and D11 (C) was performed for 30 min at 50°C at concentrations ranging from 1 to 30 μg/mL, taking only the linear regions of the reactions as initial rates. Experiments were performed in triplicate.

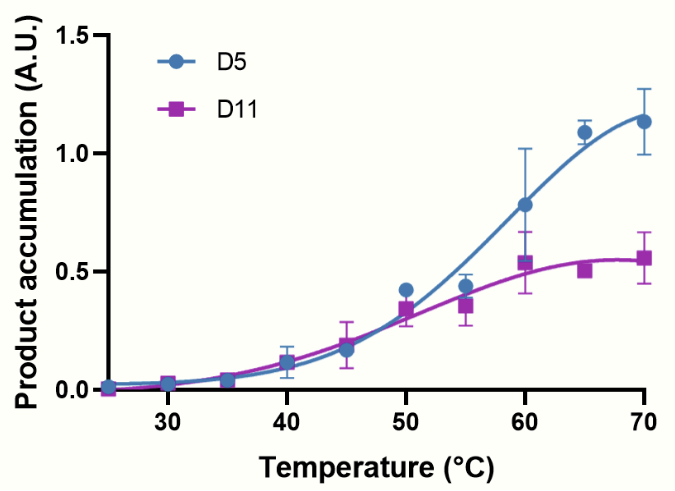


**Figure S9. Temperature ramp of PET hydrolysis by native redesigns D5 and D11.** Relative hydrolysis rates of PET films after 30 min of incubation with native redesigns D5 (A) and D11 (B), measured by absorbance at 240 nm on a multimode plate reader. Experiments were performed in triplicate.

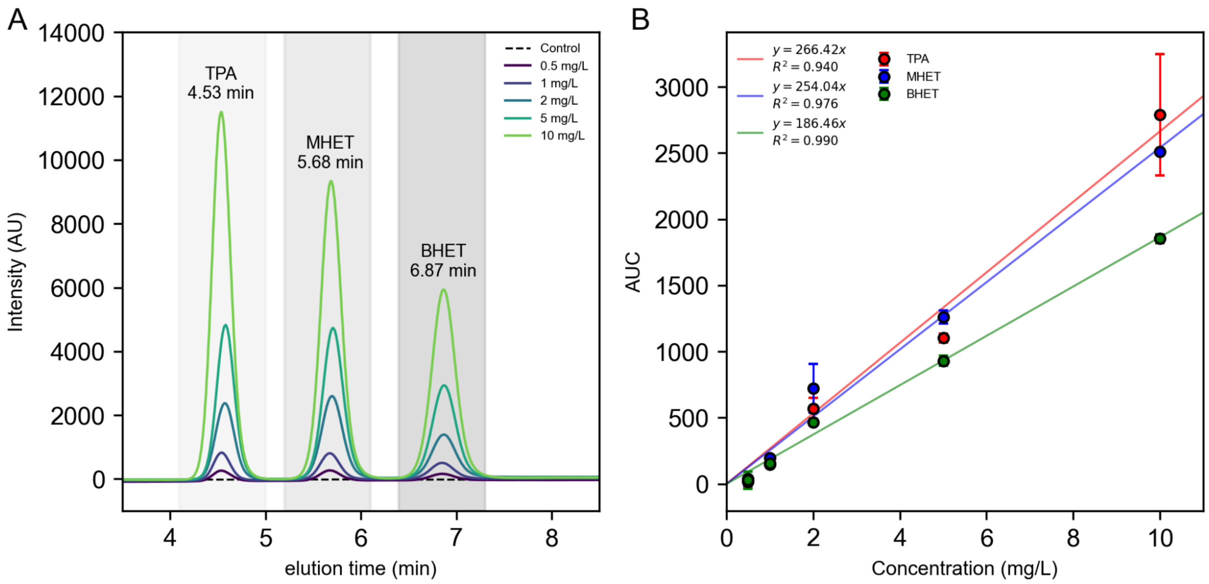


**Figure S10. Calibration curve for TPA, MHET, and BHET by HPLC.** (A) Elution profiles from HPLC runs of TPA, MHET, and BHET products to generate a calibration curve, using concentrations of mixtures of TPA, MHET, and BHET ranging from 0.5-10 mg/L of each product. A volume of 10 µL of the calibration standards were injected into the HPLC, run at a flow rate of 1.2 ml/min, and measured by absorbance at 260 nm in three independent triplicates, with the average curve for all concentrations being shown. (B) Calibration curves obtained from the analysis of the area under the curve (AUC) of the peaks for TPA, MHET, and BHET ran at different concentrations by HPLC. The whiskers correspond to the standard deviation from three independent HPLC runs.

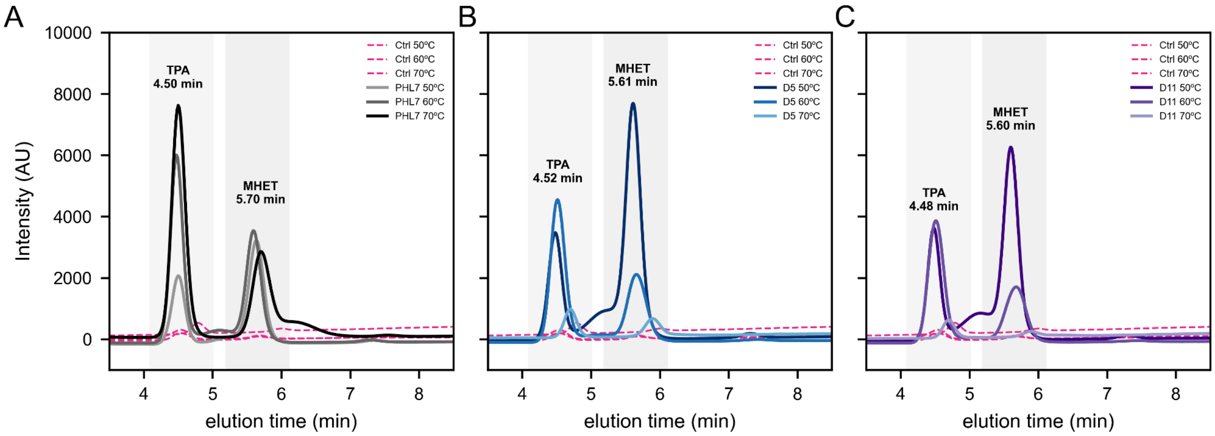


**Figure S11. Elution profiles of microPET degradation products by HPLC.** (A) Elution profiles of microPET degradation products after incubation with PHL7 for 24 h at 50°C, 60°C, and 70°C. (B) Elution profiles of microPET degradation products after incubation with design D5 for 24 h at 50°C, 60°C, and 70°C. (C) Elution profiles of microPET degradation products after incubation with design D11 for 24 h at 50°C, 60°C, and 70°C. Samples were diluted 20 times before injection of 10 µL into the HPLC. The degradation products were measured at 260 nm, and the HPLC was run at a flow rate of 1.2 ml/min. The curves correspond to the average of three independent replicates.

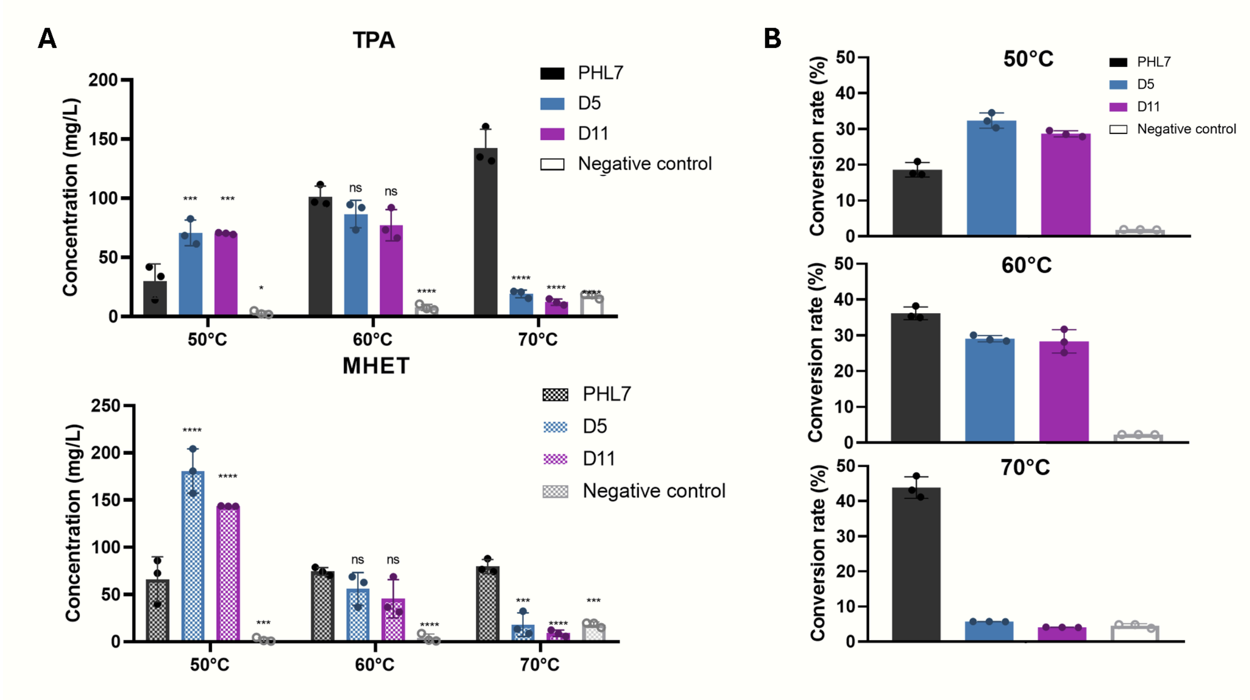


**Figure S12. Enzymatic degradation of microPET by PHL7 and active redesigns D5 and D11**. (A) Quantification of the amounts of TPA (upper bar graph) and MHET (lower bar graph) released by PHL7 and native redesigns D5 and D11 after 24 h reaction at 50, 60, and 70°C, measured in mg/L by HPLC. Product concentrations were estimated based on the calibration curves in Supplementary Figure 10. Negative controls correspond to reactions without enzyme. (B) Conversion rate (%) for PHL7 and native redesigns D5 and D11 after 24 h reaction at 50, 60, and 70°C.

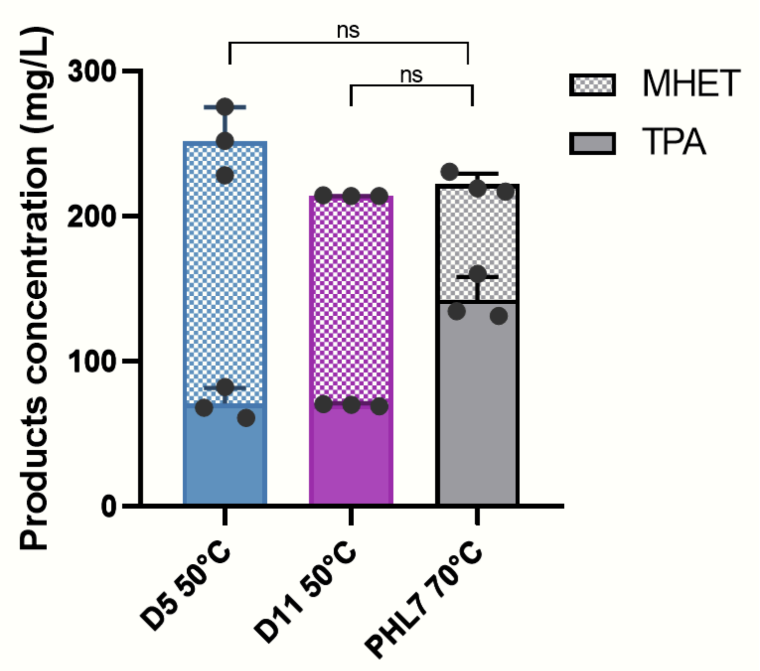


**Figure S13. Enzymatic degradation of microPET by PHL7 and the native redesigns D5 and D11 at their optimal temperatures**. Quantification in mg/L of the amounts of TPA (grey) and MHET (hatching) released by PHL7 and redesigns D5 and D11, released after 24 h microPET degradation assays at their optimal temperatures of reaction, 50ºC for D5 and D11 and 70ºC for PHL7.

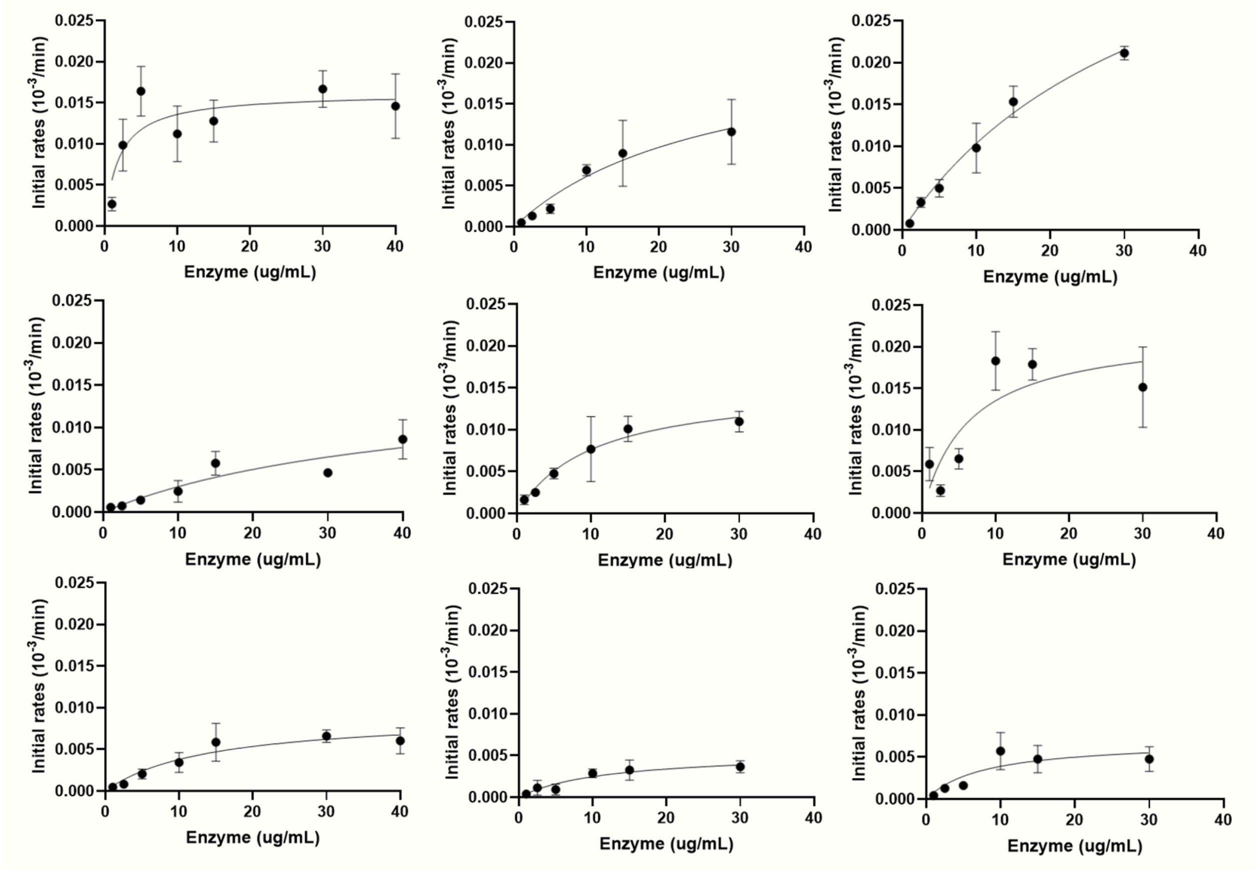


**Figure S14. Initial PET hydrolysis rates by PHL7 and native redesigns D5 and D11 at varying enzyme concentrations.** Enzymatic PET film hydrolysis was performed for 20 min at 50, 60, and 70°C at concentrations ranging 1-40 µg/mL with PHL7 (A), D5 (B), and D11 (C), determining their initial rates of degradation by collecting samples every minute for up to 20 min and measuring the degradation products via absorbance at 240 nm on a multimode plate reader. Experiments were performed in triplicate.

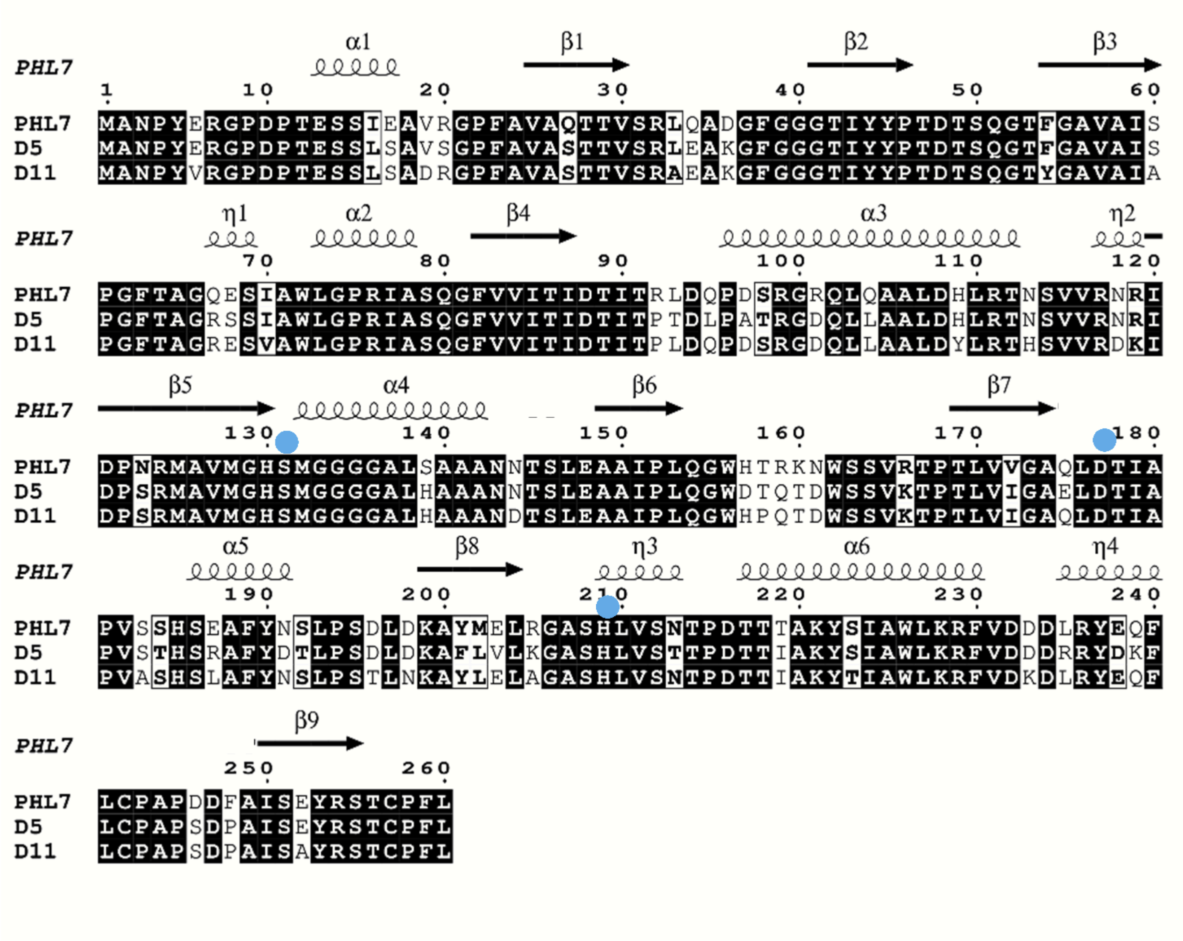

**Figure S15. Sequence variability between PHL7 and the two active native redesigns, D5 and D11**. Sequence alignment of PHL7, D5 and D11, showing the regions of strict sequence conservation between all enzymes in black background and of 70% sequence conservation in bold characters. The secondary structure topology of PHL7 (PDB 8BRB) is indicated above the alignment, and the blue dots indicate the residues from the catalytic triad.

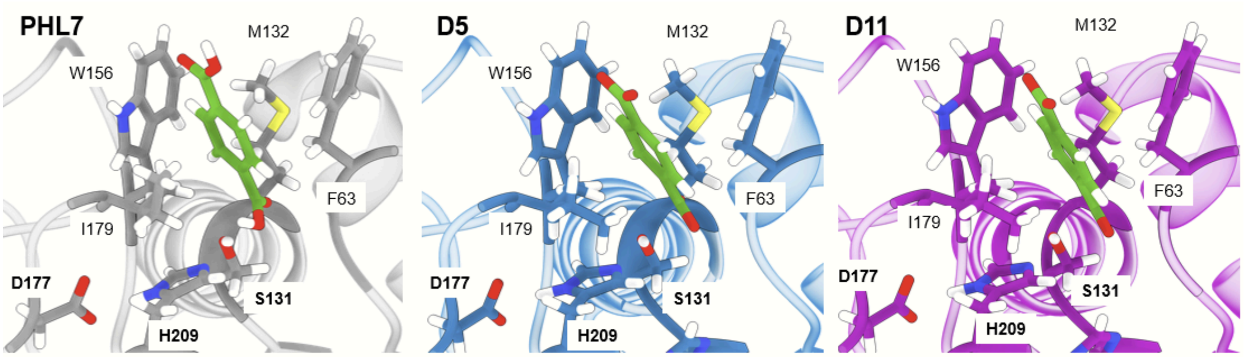


| | Catalytic triad | | | Subsite I | | | | | | | | Subsite II | | | | |
|---|---|---|---|---|---|---|---|---|---|---|---|---|---|---|---|---|
| | 131 | 177 | 209 | 63 | 132 | 156 | 179 | 93 | 95 | 185 | 189 | 64 | 65 | 69 | 130 | 210 |
| **PHL7** | S | D | H | F | M | W | I | L | Q | H | F | T | A | S | H | L |
| **D5** | S | D | H | F | M | W | I | T | L | H | F | T | A | S | H | L |
| **D11** | S | D | H | F | M | W | I | L | Q | H | F | T | A | S | H | L |

**Figure S16. Active site residues relevant for PET binding and depolymerization.** (A) Cartoon and stick representation of the experimental structure and relevant active site residues of TPA-bound PHL7 (PDB ID: 8BRB, gray) and the Boltz-2 predicted structures of D5 (blue, complex pLDDT = 0.96) and D11 (pink, complex pLDDT = 0.96) in complex with TPA. The TPA ligand is shown in green sticks. (B) MSA highlighting the variations in sequence of active site residues related to the catalytic activity of PHL7 and its binding to the TPA rings of the PET moiety (subsite I and II). Amino acids are colored by their chemistry: CNQST, green; AGILPMV, orange; RHK, red; DE, blue; FWY, yellow.

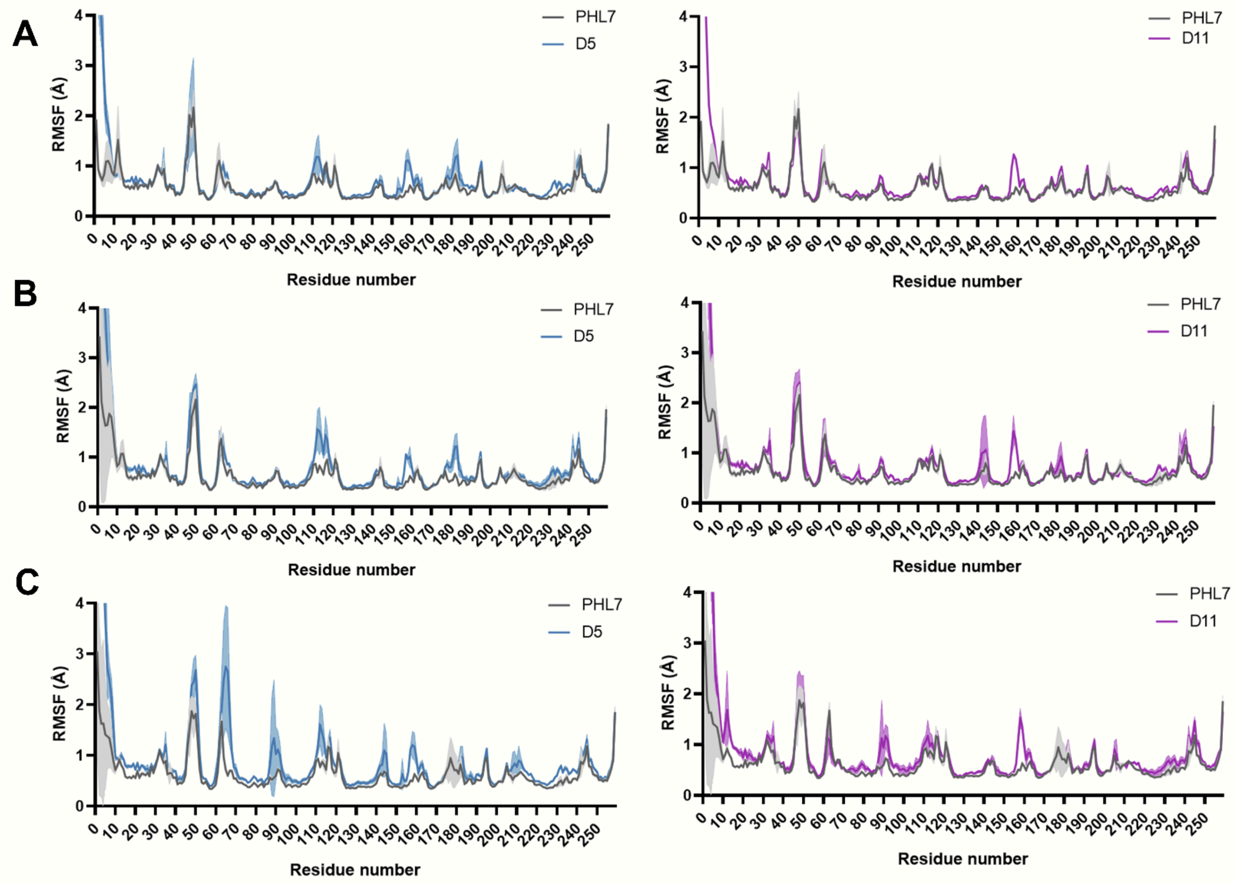


**Figure S17. Global protein flexibility of PHL7 and active designs D5 and D11 at different temperatures**. (A-C) Changes in the Cα root-mean-square fluctuations (RMSF, Å) during MD simulations of PHL7 (black), D5 (blue), and D11 (purple) at 50°C (A), 60°C (B), and 70°C (C). The shaded area represents the standard deviation over 3 × 200 ns independent trajectories per simulation system. Residue number starts with the second amino acid of each sequence (Figure S15).

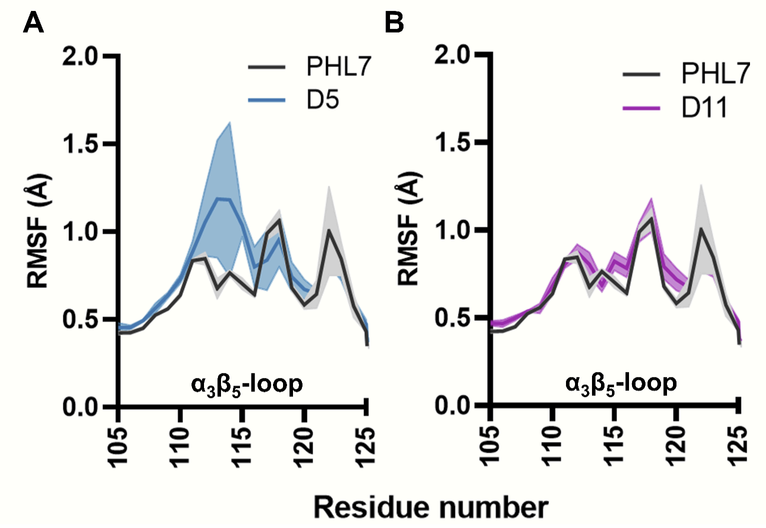


**Figure S18. Impact of residue substitutions in active designs D5 and D11 on local protein flexibility in loops outside the active site.** (A-B) Cα root-mean-square fluctuations (RMSF, Å) of the flexible $\alpha_3\beta_5$ loop (residues 105-125) during MD simulations of D5 (A) and D11 (B) designs compared to PHL7. The shaded area represents the standard deviation over 3 × 200 ns independent trajectories per simulation system. Residue number starts with the second amino acid of each sequence (Figure S15).

**Table S1. Fixed positions added into the sequence redesign process.** The group indicates the initial MSA used for the search of homologous sequences (PET PAZY or PET selected), the e-value used by hhblits to retrieve the homologous sequences (format e4/e10/e30/e50: cutoff e-value used to run hhblits; filter e4/e10/e30/e50: e-value after MAC realignment by hhblits), and the percentage of sequence conservation cutoff per position in the MSA (30, 50, 70). The group "active site" corresponds to a design campaign in which only the active site residues were fixed during the design process. For all other groups, the conserved positions of the active site were also fixed.

| Group | Fixed residues |
|---|---|
| Active site | 61 62 63 64 95 98 129 130 131 132 155 175 176 177 178 179 184 207 208 209 241 256 |
| PET selected format e4 70 | 3 32 45 60 61 80 90 92 128 130 132 133 167 173 176 208 |
| PET selected format e4 50 | 3 4 7 32 45 47 60 61 76 80 83 87 90 92 101 120 128 130 132 133 136 140 167 169 173 176 180 195 205 207 208 225 226 233 235 250 252 257 |
| PET selected format e4 30 | 2 3 4 6 7 8 9 11 12 15 16 17 22 29 30 32 34 35 40 41 43 44 45 47 54 56 57 58 59 60 61 62 73 76 77 80 81 83 87 89 90 92 101 104 105 106 107 109 118 120 123 124 126 128 129 130 132 133 136 137 139 140 143 145 146 148 149 150 151 154 164 167 169 171 172 173 176 179 180 188 191 192 194 195 197 200 204 205 207 208 222 225 226 230 233 234 235 236 239 242 249 250 252 253 256 257 |
| PET selected format e10 70 | 20 36 45 60 61 80 90 128 130 132 133 167 173 176 20 |
| PET selected format e10 50 | 20 21 22 36 38 45 52 60 61 63 76 80 83 87 90 95 101 120 128 130 132 133 136 140 144 154 158 167 169 173 176 180 205 208 214 225 229 230 257 |
| PET selected format e10 30 | 12 15 20 21 22 36 37 38 40 42 43 45 47 51 52 56 57 60 61 63 64 72 76 77 80 81 83 85 87 89 90 95 101 104 105 106 107 109 112 113 120 121 123 124 125 126 128 129 130 132 133 136 137 139 140 144 145 146 148 149 150 154 158 164 167 169 171 173 176 179 180 191 192 193 199 204 205 206 207 208 212 214 215 223 224 225 229 230 235 249 250 253 256 257 |
| PET selected format e30 70 | 40 45 54 60 61 80 92 128 130 132 133 158 167 173 176 208 256 257 |
| PET selected format e30 50 | 8 20 21 40 45 54 60 61 64 76 77 80 83 87 90 92 101 120 123 128 130 132 133 136 140 144 158 167 169 173 176 180 205 208 230 256 257 |
| PET selected format e30 30 | 2 8 10 19 20 21 22 24 40 42 43 45 51 52 54 56 57 60 61 63 64 72 73 76 77 80 81 83 85 87 89 92 93 101 104 105 106 107 109 111 113 114 115 117 119 120 121 123 124 125 126 128 129 130 132 133 136 137 140 141 144 145 146 148 150 154 158 161 164 167 169 171 172 173 176 179 180 192 193 195 197 198 200 204 205 206 207 208 225 226 230 240 250 252 253 254 255 256 257 260 |
| PET selected format e50 70 | 10 36 45 54 60 61 80 128 130 132 133 154 167 173 176 208 256 |

| | |
|---|---|
| PET selected format e50 50 | 10 20 21 36 45 54 56 60 61 64 76 77 80 83 87 101 109 112 120 123 128 130 132 133 136 140 144 154 167 169 173 176 180 194 205 208 226 230 250 256 257 260 |
| PET selected format e50 30 | 7 8 10 13 20 21 22 24 29 36 40 42 43 45 51 52 54 56 57 60 61 64 72 73 76 77 80 81 83 87 101 104 105 106 107 109 112 114 115 117 119 120 121 123 124 126 128 129 130 132 133 136 137 139 140 144 145 146 148 150 154 164 167 169 171 172 173 176 178 179 180 188 191 192 194 197 200 204 205 206 207 208 225 226 230 240 249 250 252 254 256 257 258 259 260 |
| PET selected filter e4 70 | 2 3 7 8 10 11 14 17 45 60 61 80 90 92 128 130 132 133 167 173 176 208 250 252 256 257 260 |
| PET selected filter e4 50 | 1 2 3 4 6 7 8 9 10 11 13 14 15 17 20 22 35 45 49 54 60 61 63 76 80 83 87 90 92 97 101 120 128 130 132 133 136 140 144 155 167 169 173 176 181 205 207 208 225 226 242 249 250 252 253 256 257 260 |
| PET selected filter e4 30 | 1 2 3 4 5 6 7 8 9 10 11 12 13 14 15 16 17 19 20 21 22 23 24 25 27 29 30 35 42 43 45 47 49 54 56 57 59 60 61 63 64 69 71 73 76 77 80 81 83 85 87 89 92 94 96 97 101 104 105 107 109 110 113 117 118 120 121 123 124 126 128 129 130 132 133 136 137 139 140 144 149 150 151 155 161 164 167 169 170 171 172 173 176 180 181 188 191 192 193 197 200 204 205 206 207 208 211 225 226 230 242 249 250 251 252 253 256 257 258 259 260 |
| PET selected filter e10 70 | 10 45 61 80 90 128 130 132 133 167 173 176 208 225 233 235 |
| PET selected filter e10 50 | 10 20 36 45 54 61 64 76 80 83 87 89 90 95 109 112 128 130 132 133 136 139 144 167 169 173 176 180 205 207 208 214 225 233 234 235 236 239 240 249 250 |
| PET selected filter e10 30 | 4 9 10 11 20 24 28 29 34 36 43 44 45 51 54 56 57 58 61 64 65 72 76 77 80 81 83 85 87 89 95 104 105 106 107 109 111 112 113 114 115 117 119 123 124 126 128 129 130 131 133 134 137 138 139 140 141 145 147 149 150 151 152 153 155 162 165 168 170 171 172 173 174 177 180 181 189 190 192 193 194 198 200 203 205 206 207 208 209 215 226 227 230 231 233 234 235 236 237 240 241 243 250 251 255 |
| PET selected filter e30 70 | 7 10 20 21 44 45 61 77 78 80 82 83 113 120 128 129 130 132 133 134 139 154 167 173 176 189 193 205 208 225 230 233 250 256 257 |
| PET selected filter e30 50 | 2 3 4 6 7 8 10 11 14 20 21 22 24 42 43 44 45 51 54 61 76 77 78 79 80 81 82 83 88 102 105 109 113 114 117 120 123 125 126 128 129 130 131 132 133 134 135 136 139 140 144 148 152 153 154 167 168 169 173 176 179 180 181 189 192 193 197 198 202 205 208 224 225 226 230 233 250 252 256 257 259 260 |
| PET selected filter e30 30 | 2 3 4 5 6 7 8 9 10 11 12 13 14 16 17 19 20 21 22 23 24 26 28 29 34 36 39 40 41 42 43 44 45 46 51 52 54 55 56 57 59 60 61 62 64 67 69 71 72 73 74 75 76 77 78 79 80 81 82 83 84 85 86 88 89 93 95 98 99 100 102 103 104 105 106 107 108 109 111 112 113 114 115 116 117 118 119 120 121 123 125 126 127 128 129 130 131 132 133 134 135 136 137 138 139 140 141 143 144 145 146 147 148 149 150 151 152 153 154 155 162 164 166 167 168 169 170 171 172 173 175 176 177 178 179 180 181 182 184 187 189 190 192 193 196 197 198 200 202 203 205 206 207 208 211 215 221 223 224 225 226 227 229 230 232 233 235 236 240 242 246 248 249 250 252 253 254 255 256 257 259 260 |
| PET selected filter e50 70 | 2 3 7 8 10 11 14 17 20 21 22 29 36 39 41 42 43 44 45 46 54 57 58 60 61 71 73 74 76 77 78 80 81 82 83 86 88 89 93 95 98 102 104 105 106 109 113 120 123 126 128 130 131 132 133 134 135 137 139 140 146 147 148 152 154 155 164 167 168 169 170 176 179 189 193 198 200 202 208 225 227 229 231 233 235 236 239 240 250 256 257 258 |

| | |
|---|---|
| PET selected filter e50 50 | 2 3 4 6 7 8 9 10 11 13 14 17 20 21 22 24 26 28 29 36 37 38 39 40 41 42 43 44 45 46 50 51 52 54 55 56 57 58 60 61 62 64 67 68 71 73 74 75 76 77 78 79 80 81 82 83 84 85 86 87 88 89 92 93 94 97 98 100 101 104 105 106 107 109 110 113 114 119 120 123 124 125 126 127 128 129 130 131 132 133 134 135 137 139 140 144 145 146 147 148 149 150 151 152 153 154 155 162 164 166 167 168 169 170 174 175 176 177 179 180 183 184 185 187 188 189 190 191 192 193 196 198 199 200 202 205 206 208 211 212 213 215 219 223 225 226 227 228 229 231 233 234 235 236 238 239 240 241 242 249 250 252 253 254 255 256 257 259 260 |
| PET selected filter e50 30 | 1 2 3 4 5 6 7 8 9 10 11 12 13 14 15 16 17 18 19 20 21 22 23 24 25 26 28 29 30 31 32 34 35 36 37 38 39 40 41 42 43 44 45 46 47 48 49 50 51 52 53 54 55 56 57 58 59 60 61 62 63 64 65 66 67 68 69 70 71 72 73 74 75 76 77 78 79 80 81 82 83 84 85 86 87 88 89 90 91 92 93 94 95 96 97 98 99 100 101 102 103 104 105 106 107 108 109 110 111 112 113 114 115 116 117 118 119 120 121 122 123 124 125 126 127 128 129 130 131 132 133 134 135 136 137 138 139 140 141 143 144 145 146 147 148 149 150 151 152 153 154 155 156 157 159 160 161 162 163 164 166 167 168 169 170 171 172 173 174 175 176 177 178 179 180 181 182 183 184 185 186 187 188 189 190 191 192 193 194 195 196 197 198 199 200 201 202 203 204 205 206 207 208 209 210 211 212 213 214 215 216 217 219 220 221 222 223 225 226 227 228 229 230 231 232 233 234 235 236 237 238 239 240 241 242 243 244 246 247 248 249 250 252 253 254 255 256 257 258 259 260 |
| PET PAZY format e4 70 | 2 3 4 6 7 8 10 11 20 35 45 60 61 80 91 128 130 132 133 167 173 176 208 256 258 |
| PET PAZY format e4 50 | 2 3 4 5 6 7 8 9 10 17 20 21 25 34 35 45 54 60 61 63 76 80 83 87 89 91 95 97 101 117 120 128 130 132 133 136 140 144 154 167 169 173 176 180 205 207 208 225 226 250 252 254 256 258 259 260 |
| PET PAZY format e4 30 | 1 2 3 4 5 6 7 8 9 10 13 14 15 17 20 21 25 27 34 35 38 40 42 43 45 51 52 54 56 57 58 60 61 63 64 69 73 76 77 80 81 82 83 87 88 89 90 91 95 97 101 104 106 107 109 111 114 117 119 120 121 123 124 126 128 129 130 132 133 136 137 139 140 144 145 146 148 149 150 151 154 158 164 167 169 171 172 173 176 179 180 188 191 192 193 197 200 204 205 206 207 213 222 225 226 230 240 249 250 252 253 254 255 256 257 258 259 260 |
| PET PAZY format e10 70 | 3 35 36 60 61 80 90 128 130 132 167 173 176 208 256 |
| PET PAZY format e10 50 | 3 35 36 54 60 61 63 76 80 83 87 89 90 95 101 120 128 130 132 133 136 140 144 152 167 169 173 176 180 205 208 211 225 226 256 |
| PET PAZY format e10 30 | 2 3 4 8 10 14 15 17 21 31 35 36 42 44 51 52 54 56 57 58 60 61 63 64 71 72 73 76 77 80 81 83 87 89 90 95 101 104 105 107 109 117 120 121 123 124 126 128 129 130 132 133 134 136 137 139 140 144 145 146 148 149 150 151 152 157 161 164 167 169 170 171 172 173 176 179 180 188 191 192 193 200 204 205 206 207 208 211 215 216 222 225 226 230 239 240 250 252 253 254 255 256 257 258 |
| PET PAZY format e30 70 | 3 21 45 54 60 61 80 120 128 130 132 133 167 173 176 208 256 |
| PET PAZY format e30 50 | 3 7 8 20 21 24 45 54 60 61 63 76 77 80 83 87 101 109 120 123 128 130 132 133 136 140 144 158 167 169 173 176 180 205 208 214 225 226 230 250 256 |
| PET PAZY format e30 30 | 2 3 4 6 7 8 9 10 11 16 19 20 21 22 24 40 42 43 45 51 52 54 56 57 60 61 63 64 74 75 76 77 80 81 83 85 87 88 91 95 101 104 105 106 107 109 115 117 119 120 121 123 124 125 126 128 129 130 132 133 136 137 139 140 141 144 145 146 148 150 151 154 157 158 164 167 169 171 172 173 176 179 180 187 189 192 195 196 197 199 202 204 205 206 207 208 214 225 226 230 240 246 250 252 253 254 255 256 257 259 260 |

| | |
|---|---|
| PET PAZY format e50 70 | 3 10 45 54 60 61 80 90 92 128 130 132 133 167 173 176 208 227 241 |
| PET PAZY format e50 50 | 3 8 10 20 21 22 45 54 60 61 63 76 77 80 83 87 89 90 92 95 96 101 109 120 123 128 130 132 133 136 140 144 154 167 169 173 176 179 205 208 223 227 229 240 241 253 256 |
| PET PAZY format e50 30 | 2 3 4 6 7 8 10 11 13 20 21 22 24 34 39 42 43 45 49 51 52 54 56 57 58 60 61 63 68 73 76 77 80 81 83 87 89 90 92 94 95 96 101 104 105 106 107 109 111 117 119 120 121 123 124 126 128 129 130 132 133 136 137 139 140 141 144 145 154 155 161 164 167 169 171 172 173 176 178 179 188 191 192 193 197 200 204 205 206 207 208 214 217 219 220 221 222 223 225 227 228 229 230 231 232 236 239 240 241 246 252 253 256 |
| PET PAZY filter e4 70 | 3 4 6 7 8 10 36 45 54 60 61 80 101 120 128 130 132 133 167 173 176 208 256 |
| PET PAZY filter e4 50 | 2 3 4 6 7 8 9 10 11 13 14 25 35 36 45 54 60 61 64 76 77 80 83 88 101 105 120 123 128 130 132 133 136 140 148 154 167 169 172 173 176 179 180 192 196 204 205 208 230 250 256 |
| PET PAZY filter e4 30 | 2 3 4 5 6 7 8 9 10 11 12 13 14 17 25 27 35 36 38 40 42 43 45 50 51 52 54 56 57 60 61 64 76 77 80 81 82 83 84 86 87 88 101 104 105 107 109 111 113 114 115 117 119 120 121 123 124 125 126 128 130 132 133 135 136 137 139 140 143 145 146 147 148 150 153 154 167 169 170 172 173 176 179 180 185 188 191 192 195 196 198 199 200 203 204 205 207 208 209 225 226 230 240 250 252 253 255 256 257 258 259 |
| PET PAZY filter e10 70 | 3 10 35 45 54 60 61 80 101 120 128 130 132 133 167 173 176 208 250 256 |
| PET PAZY filter e10 50 | 2 3 4 6 7 8 10 11 20 27 34 35 45 54 60 61 62 63 64 76 77 80 83 101 105 120 121 123 128 130 132 133 136 140 148 158 167 169 173 176 179 180 189 204 205 208 230 250 252 256 |
| PET PAZY filter e10 30 | 1 2 3 4 6 7 8 9 10 11 13 14 15 17 20 21 27 29 31 34 35 38 40 42 43 45 51 52 54 56 57 60 61 62 63 64 69 73 76 77 80 81 82 83 87 89 91 93 94 95 97 101 104 105 107 109 113 114 117 119 120 121 123 124 125 126 128 130 131 132 133 135 136 137 140 141 143 144 145 148 150 153 158 162 165 167 169 170 172 173 176 179 180 185 189 192 193 198 199 200 203 204 205 207 208 209 214 225 226 230 240 242 250 252 253 254 256 257 259 260 |
| PET PAZY filter e30 70 | 3 6 7 8 10 11 20 21 36 45 54 61 77 78 80 83 113 120 128 130 132 133 143 144 167 173 176 208 230 252 256 258 259 260 |
| PET PAZY filter e30 50 | 2 3 4 6 7 8 9 10 11 14 20 21 22 24 36 41 42 43 44 45 51 54 61 76 77 78 79 80 81 82 83 85 102 107 113 117 120 123 125 126 128 129 130 132 133 136 137 140 143 144 145 148 161 167 169 172 173 176 180 189 197 207 208 215 224 225 226 230 231 232 233 239 249 250 252 253 256 257 258 259 260 |
| PET PAZY filter e30 30 | 2 3 4 5 6 7 8 9 10 11 13 14 17 18 20 21 22 24 25 28 31 32 36 38 41 42 43 44 45 46 47 48 51 52 53 54 55 56 57 58 59 60 61 63 69 72 73 75 76 77 78 79 80 81 82 83 84 85 86 87 90 93 98 101 102 104 105 106 107 108 109 110 112 113 114 115 116 117 118 119 120 121 123 124 125 126 128 129 130 131 132 133 134 136 137 139 140 142 143 144 145 146 147 148 150 152 153 154 157 161 166 167 168 169 170 172 173 176 178 179 180 181 182 184 185 188 189 192 193 197 198 203 205 206 207 208 209 210 211 215 216 222 224 225 226 227 228 230 231 232 233 236 239 240 241 246 249 250 251 252 253 254 255 256 257 258 259 260 |
| PET PAZY filter e50 70 | 2 3 6 7 8 10 11 14 17 20 21 22 29 36 39 41 42 43 44 45 46 54 57 60 61 64 71 73 74 77 78 80 81 82 83 86 88 89 93 95 98 102 104 105 106 109 113 120 123 126 128 130 131 132 133 134 135 137 139 140 144 145 146 147 148 149 152 154 155 167 168 169 170 176 179 180 184 189 193 196 198 200 202 208 225 227 229 231 233 235 236 239 240 250 256 258 259 260 |

| | |
|---|---|
| PET PAZY filter e50 50 | 2 3 4 6 7 8 9 10 11 13 14 17 19 20 21 22 24 28 29 30 36 37 38 39 40 41 42 43 44 45 46 48 50 51 52 54 55 56 57 58 60 61 62 63 64 67 68 70 71 72 73 74 75 76 77 78 79 80 81 82 83 84 85 86 87 88 89 92 93 94 95 96 97 98 99 101 102 104 105 106 107 109 110 113 114 115 116 119 120 123 124 125 126 127 128 129 130 131 132 133 134 135 137 139 140 144 145 146 147 148 149 150 151 152 153 154 155 162 164 166 167 168 169 170 174 175 176 177 179 180 181 183 184 185 187 188 189 190 191 192 193 196 198 199 200 202 205 206 208 211 212 213 215 219 223 225 226 227 228 229 231 233 234 235 236 238 239 240 241 242 249 250 252 253 254 255 256 257 258 259 260 |
| PET PAZY filter e50 30 | 1 2 3 4 5 6 7 8 9 10 11 12 13 14 15 16 17 18 19 20 21 22 23 24 26 29 30 31 33 35 36 37 38 39 40 41 42 43 44 45 46 47 48 49 50 51 52 53 54 55 56 57 58 59 60 61 62 63 64 66 67 68 69 70 71 72 73 74 75 76 77 78 79 80 81 82 83 84 85 86 87 88 89 92 93 94 95 96 97 98 99 100 101 102 103 104 105 106 107 108 109 110 111 112 113 114 115 116 117 118 119 120 121 122 123 124 125 126 127 128 129 130 131 132 133 134 135 136 137 139 140 143 144 145 146 147 148 149 150 151 152 153 154 156 157 158 159 160 161 162 163 164 165 166 168 169 170 171 172 173 174 175 176 177 178 179 180 181 182 183 184 185 186 187 188 189 190 191 192 193 194 195 196 197 198 199 200 201 202 203 204 205 206 207 208 209 210 211 212 213 214 215 216 217 218 219 221 222 223 224 225 226 227 228 229 230 231 232 233 234 235 236 238 239 240 241 242 244 249 250 251 252 253 254 255 256 257 258 259 260 261 |

**Table S2. Selected designed sequences for experimental screening.** These sequences were selected using a Levenshtein matrix that clustered all 1176 sequences in 36 groups. The group indicates the initial MSA used for the search of homologous sequences (PET PAZY or PET selected), the e-value used by hhblits to retrieve the homologous sequences (format e4/e10/e30/e50: cutoff e-value used to run hhblits; filter e4/e10/e30/e50: e-value after MAC realignment by hhblits), and the percentage of sequence conservation cutoff per position in the MSA (30, 50, 70). The sequence number indicates which of the 12 sequences generated by either ProteinMPNN or LigandMPNN was selected. The sequence identity percentage of all designed sequences was calculated against PHL7.

| Name | Group | Sequence number | Deep learning tool | Amino acid sequence | Sequence Identity (%) |
|---|---|---|---|---|---|
| PHL7 | - | - | - | MANPYERGPDPTESSIEAVRGPFAVA<br>QTTVSRLQADGFGGGTIYYPTDTSQG<br>TFGAVAISPGFTAGQESIAWLGPRIA<br>SQGFVVITIDTITRLDQPDSRGRQLQ<br>AALDHLRTNSVVRNRIDPNRMAVMGH<br>SMGGGGALSAAANNTSLEAAIPLQGW<br>HTRKNWSSVRTPTLVVGAQLDTIAPV<br>SSHSEAFYNSLPSDLDKAYMELRGAS<br>HLVSNTPDTTTAKYSIAWLKRFVDDD<br>LRYEQFLCPAPDDFAISEYRSTCPFL | 100 |
| D1 | PET PAZY filter e10 30 | 3 | LigandMPNN | MANPYVRGPDPTESSISAAAGPFATA<br>STTVTRAEADGFGGGTIYYPTDSSQG<br>TYGAVAIAPGFTAGRDSIAWLGPAIA<br>SQGFVVIVIDTITRTDQPASRGDQLL<br>AALDYLKTNSVVKNLIDPSRMAVMGH<br>SMGGGGALHAAANNTSLKAAIPLQGW<br>DPRTDWSSIRTPTLVIGAELDTIAPV<br>ETHSLAFYNTLPSDLNKAYLELRGAS<br>HLVATTPNTTILRYTVAWLRLFVDDD<br>RRYEKFLCPAPSDPAISAYRSTCPFL | 78.38 |
| D2 | PET PAZY filter e10 30 | 7 | LigandMPNN | MANPYVRGPDPTESSISAASGPFAVA<br>STTVSRAAADGFGGGTIYYPTDTSQG<br>TYGAVAIAPGFTAGRDSIEWWGPAIA<br>SKGFVVIVIDTITRTDQPASRGDQLL<br>AALDYLKTNSVVKNKIDPSRMAVMGH<br>SMGGGGALHAAANNTSLKAAIPLQGW<br>DPRTDWSSIRTPTLVIGAELDTIAPV<br>ATHSLAFYNSLPSTLNKAYLELRGAS<br>HLVATTPNTTILKYTVAWLRLFVDND<br>RRYEKFLCPAPSDPAISAYRSTCPFL | 78.38 |
| D3 | PET PAZY filter e10 50 | 5 | LigandMPNN | MANPYVRGPDPTESSLSAATGPFATA<br>STEVSAAEADGFGGGTIYYPTDTSQG<br>KYGAVAIAPGFTAGRDSVEWLGPAIA<br>SKGFVVIVIDTTTLTDGPATRGDQLL<br>AALDYLKTHPAVKDKIDPSRRAVMGH<br>SMGGGGALHAAANDSSLKATIPLVGW<br>DPQTDWSSVKTPTLVIGAELDTIAPV<br>ETHSRAFYDTLPSDLNKAYLELRGAS<br>HLVATTPNETILKYTVAWLRLFVDDD<br>RRYEKFLCPAPSDPAISAYDSTCPFK | 69.88 |

| | | | | | |
|---|---|---|---|---|---|
| D4 | PET PAZY filter e30 30 | 4 | ProteinMPNN | MANPYERGPDPTESSLAAVTGPFAVA STTVSRLEAKGFGGGTIYYPTDTSQG TFGAVAISPGFTAGRSSIAWLGPRIA SQGFVVITIDHLTPTDDPETRGKQLL AALDHLRTNSVVRNRIDPSRMAVMGH SMGGGGALYAAANNTSLEAVIPLQGW STRKDWSSIRTPTLVIGAELDTIAPV STHSRAFYDSLPADLDKAYLVLKGAS HLVSTSPDTTILRYSIAWLKRFVDDD ERYDRFLCPAPTDPRISEYRSTCPFL | 83.01 |
| D5 | PET PAZY filter e30 30 | 6 | LigandMPNN | MANPYERGPDPTESSLSAVSGPFAVA STTVSRLEAKGFGGGTIYYPTDTSQG TFGAVAISPGFTAGRSSIAWLGPRIA SQGFVVITIDTITPTDLPATRGDQLL AALDHLRTNSVVRNRIDPSRMAVMGH SMGGGGALHAAANNTSLEAAIPLQGW DTQTDWSSVKTPTLVIGAELDTIAPV STHSRAFYDTLPSDLDKAFLVLKGAS HLVSTTPDTTIAKYSIAWLKRFVDDD RRYDKFLCPAPSDPAISEYRSTCPFL | 84.56 |
| D6 | PET PAZY filter e30 30 | 6 | ProteinMPNN | MANPYERGPDPTLSSLEAVTGPFAVA STTVSRLEAKGFGGGTIYYPTDTSQG TFGAVAISPGFTATRSSIDWLGPRIA SQGFVVITIDHLTLTDGPEERGKQLL AALDHLRTNSVVRNRIDPSRMAVMGH SMGGGGALYAAANNTSLEAVIPLQGW STKTDWSSIKTPTLVIGAELDTIAPV STHSRKFYDTLPADLDKAFLVLKGAS HLVSTSPDTTISRYSIAWLKRFVDDD RRYDRFLCPAPSDPRISEYRSTCPFL | 79.92 |
| D7 | PET PAZY filter e30 30 | 9 | ProteinMPNN | MANPYERGPDPTLSSLSAVTGPFAVA STTVSRLEAKGFGGGTIYYPTDTSQG TFGAVAISPGFTAGRSSIAWLGPRIA SQGFVVITIDSLTLTDGPEERGAQLL AALDHLRTNSVVRNRIDPSRMAVMGH SMGGGGALYAAANNTSLEAVIPLQGW STQKDWSSIKTPTLVIGAELDTIAPV STHSKAFYDSLPSDLDKAYLVLKGAS HLVSTTPDTTIARYSIAWLKRFVDDD RRYDRFLCPAPTDPAISEYRSTCPFL | 83.4 |
| D8 | PET PAZY filter e4 30 | 11 | LigandMPNN | MANPYERGPDPTESSLSAASGPFAVA STTVTAAEADGFGGGTIYYPTDTSQG TYGAVAIAPGFTAGRDSVEWLGPAIA SHGFVVIVIDTTTLTDGPATRGDQLL AALDYLKTNSVVKNKIDPSRMAVMGH SMGGGGALHAAANNSSLEATIPLQGW DPQKDWSSIKTPTLVIGAELDTIAPV ETHSRAFYDSLPSDLNKAYLELRGAS HLVATTPNETILKYTVAWLRTFVDKD RRYEKFLCPAPSDPAISAYRSTCPFL | 74.9 |

| | | | | | |
|---|---|---|---|---|---|
| D9 | PET PAZY filter e4 50 | 5 | LigandMPNN | MANPYVRGPDPTESSLSARTGPFAVA STEVTAAEADGFGGGTIYYPTDTSQG RYGAVAIAPGFTAGRSSVEWLGPAIA SHGFVVIVIDTTTLTDGPATRGDQLL AALDYLKTHPDVKDKIDPSRRAVMGH SMGGGGALHAAANDSSLKATIPLVGW DPQTDWSSIKTPTLVIGAELDTIAPV ATHSRAFYNSLPSDLNKAYLELRGAS HLVATTPNTTILRYTVAWLRTFVDDD KRYEKFLCPAPSDPAISAYDSTCPFK | 70.27 |
| D10 | PET PAZY filter e50 30 | 9 | LigandMPNN | MANPYERGPDPTESSIEAVRGPFAVD QTTVSRAQADGFGGGTIYYPTDTSQG TFGAVAISPGFTAGQESIAWLGPRIA SQGFVVITIDTITPLDQPDSRGRQLQ AALDHLRTNSVVRNRIDPNRMAVMGH SMGGGGALYAAANNTSLEAAIPLQGW HTRKNWSSVRTPTLVVGAQLDTIAPV SSHSEAFYNSLPSDLDKAYMELRGAS HLVSNTPDTTTAKYSIAWLKRFVDDD LRYEQFLCPAPSDPRISEYRSTCPFL | 96.91 |
| D11 | PET PAZY filter e50 50 | 9 | LigandMPNN | MANPYVRGPDPTESSLSADRGPFAVA STTVSRAEAKGFGGGTIYYPTDTSQG TYGAVAIAPGFTAGRESVAWLGPRIA SQGFVVITIDTITPLDQPDSRGDQLL AALDYLRTHSVVRDKIDPSRMAVMGH SMGGGGALHAAANDTSLEAAIPLQGW HPQTDWSSVKTPTLVIGAQLDTIAPV ASHSLAFYNSLPSTLNKAYLELAGAS HLVSNTPDTTIAKYTIAWLKRFVDKD LRYEQFLCPAPSDPAISAYRSTCPFL | 84.17 |
| D12 | PET PAZY filter e50 50 | 12 | ProteinMPNN | MANPYIRGPDPTESSLSAERGPFAVS STTVSAAEAVGFGGGTIYYPTDTSQG TYGAVAIAPGFTAGRESVAWLGPRIA SQGFVVITIDTISPLDQPDSRGAQLL AALDYLRTHSVVRDKIDPSRMAVMGH SMGGGGALYAAANDTSLEAAIPLQGW SPTLDWSSVKTPTLVIGAQLDTIAPV ASHSRAFYNSLPSNLNKAYLEIAGAS HLVSNTPDATIARYTIAWLKRFVDRD LRYDQFLCPAPTDPDISDYRSTCPFL | 80.69 |
| E1 | PET PAZY format e10 30 | 1 | LigandMPNN | MANPYVKGPAPTEASISAATGPFAVD STTVTRAEADGFGGGTIYYPTDTSQG TYGAVAIAPGFTAGRSSVEWLGPAIA SHGFVVIVIDTITPTDGPATRGDQLL AALDYLKTHPAVKNKIDPSRMAVMGH SMGGGGALHAAANDTSLKAAIPLVGW DTQTDWSSVKTPTLVVGAELDTIAPV ATHSLAFYNSLPSTLNKAYLELRGAS HLVSTSPDTTILRYSVAWLRTFVDDD RRYEQFLCPAPSDPAISAYRSTCPFS | 74.52 |

| | | | | | |
|---|---|---|---|---|---|
| E2 | PET PAZY format e30 30 | 6 | LigandMPNN | MANPYVRGPDPTEASLEADRGPFAVASTTVTAAEAKGFGGGTIYYPTDTSQGTYGAVAIAPGFTAGRDSVEWLGPRIASHGFVVITIDTTTRTDGPATRGDQLLAALDYLKTHPAVKNKIDPSRMAVMGHSMGGGGALHAAANDTSLKAAIPLVGWDTRTDWSSVKTPTLIVGAELDTIAPVATHSRAFYDSLPSDLDKAYLELRGASHLVATTPNTTIHKYTVAWLRRFVDNDRRYEKFLCPAPSDPRISAYRSTCPFL | 75.68 |
| E3 | PET PAZY format e30 30 | 11 | LigandMPNN | MANPYVRGPDPTRASLEADRGPFAVDSTEVTREEAKGFGGGTIYYPTDTSQGTYGAVAIAPGFTAGRDSVEWWGPRIASRGFVVITIDTITRTDGPATRGDQLLAALDYLKTHPAVKNKIDPSRMAVMGHSMGGGGALHAAANDTSLKAAIPLVGWDTRTDWSSVKTPTLIVGAELDTIAPVATHSLAFYNSLPADLDKAYLELRGASHLVATTPNTTIHRYTVAWLRRFVDGDRRYEQFLCPAPADPAISAYRSTCPFL | 75.29 |
| E4 | PET PAZY format e4 30 | 1 | LigandMPNN | MANPYERGPDPTESSISARSGPFAVASTEVTAAEADGFGGGTIYYPTDTSQGTYGAVAIAPGFTAGRDSIEWLGPAIASHGFVVIVIDTITRTDEPASRGDQLLAALDYLKTHPVVKNKIDPSRMAVMGHSMGGGGALHAAANDTSLKAAIPLVGWDPRTDWSSVKTPTLIVGAELDTIAPVATHSLAFYNSLPSTLDKAYLELRGASHLVATTPNETILRYSVAWLRRFVDNDRRYEKFLCPAPSDPAISAYRSTCPFL | 76.45 |
| E5 | PET PAZY format e4 30 | 11 | LigandMPNN | MANPYERGPDPTESSISAASGPFATASTEVTAAEADGFGGGTIYYPTDTSQGTYGAVAIAPGFTAGRDSIAWWGPAIASKGFVVIVIDTITRTDGPASRGDQLLAALDYLKTHPVVKNKIDPSRMAVMGHSMGGGGALHAAANDTSLKAAIPLVGWDPRTDWSSVKTPTLVVGAELDTIAPVATHSRAFYDSLPADLDKAYLELRGASHLVATTPNTTILRYSVAWLRRFVDDDKRYEKFLCPAPSDPAISAYRSTCPFL | 76.83 |
| E6 | PET PAZY format e50 30 | 9 | LigandMPNN | MANPYVRGPEPTESSLSAASGPFAVDSTEVTREEAKGFGGGTIYYPTDTSQGTYGAVAIAPGFTAGRDSVEWWGPRIASHGFVVIVIDTITPLDQPDVRGDQLLAALDYLKTHPAVKNKIDPSRMAVMGHSMGGGGALHAAAKDTSLKAAIPLVGWYPRTDWSSVKTPTLIVGAELDTIAPVATHSRAFYDSLPSDLDKAYLELRGASHLVATTPNTTIAKYSIAWLKRFVDDDKRYDKFLCPAPSDPAISAYRSTCPFS | 75.68 |

| | | | | | |
|---|---|---|---|---|---|
| E7 | PET selected filter e10 30 | 6 | LigandMPNN | MANPYIHGPDPTEASLSAAAGPFAVA<br>STTVTAAEAKGFGGGTIYYPTDTSQG<br>KYGAVAISPGFTAGRSSVEWLGPAIA<br>SHGFVVITIDTITLTVGPATRGAQLL<br>AALDYLKTNSVVKNKIDPSRMAVMGH<br>SMGGGGALSAAANDSSLEAAIPLQGW<br>DPQTDWSSIRTPTLVVGAQLDTIAPV<br>ATHSRAFYNTLPSTLNKAYLELAGAS<br>HLVATSPDTTILRYTVAWLKLFVDKD<br>LRYEKFLCPAPSDPAISEYDSTCPFS | 74.52 |
| E8 | PET selected filter e30 30 | 2 | LigandMPNN | MANPYERGPDPTESSLEAERGPFAVD<br>QTTVTAAEAKGFGGGTIYYPTDTSQG<br>TYGAVAISPGFTAGRESIEWLGPRIA<br>SQGFVVITIDTITLTDGPDTRGRQLQ<br>AALDHLKTNSVVRNRIDPSRRAVMGH<br>SMGGGGALSAAANNTSLEAAIPLQGW<br>YGTTDWSSVKTPTLVVGAELDTIAPV<br>STHSLAFYNSLPSTLDKAYLELAGAS<br>HLVSTTPDTTIAKYTIAWLKRFVDDD<br>KRYEKFLCPAPTDPAISAYRSTCPFL | 84.94 |
| E9 | PET selected filter e30 30 | 5 | ProteinMPNN | MANPYERGPDPTESSLEAERGPFAVD<br>QATVSAAEAVGFGGGTIYYPTDTSQG<br>TYGAVAISPGFTAGRESIAWLGPRIA<br>SQGFVVITIDTISPTDGPEARGRQLQ<br>AALDHLKTNSVVRNRIDPSRRAVMGH<br>SMGGGGALSAAAANTSLEAAIPLQGW<br>SPTRDWSSVRTPTLVVGAELDTIAPV<br>STHDLAFYNSLPSSLDKAYLELAGAS<br>HLASTTPDRTISRYTIAWLKLFVDDD<br>RRYERFLCPAPTDPAISNYRSTCPFL | 82.24 |
| E10 | PET selected filter e30 30 | 9 | ProteinMPNN | MANPYERGPDPTESSLEAERGPFAVD<br>QTTVTAAEAKGFGGGTIYYPTDLSQG<br>TYGAVAISPGFTATRESIAWLGPRIA<br>SQGFVVITIDTISLTDGPEERGRQLQ<br>AALDHLKTNSVVRNRIDPSRRAVMGH<br>SMGGGGALSAAAANTSLEAAIPLQGW<br>SPTLDWSSVRTPTLVVGAELDTIAPV<br>STHELAFYNSLPSDLDKAYLELKGAS<br>HLASTSPDRTIARYSIAWLKLFVDDD<br>RRYERFLCPAPTDPAISNYRSTCPFL | 82.24 |
| E11 | PET selected filter e4 30 | 5 | LigandMPNN | MANPYERGPDPTESSIEAERGPFAVA<br>STTVSAAAADGFGGGTIYYPTDTSQG<br>KYGAVAISPGFTAGRDSIEWLGPAIA<br>SHGFVVITIDTISLLDQPDSRGDQLL<br>AALDYLRTNSTVKNRIDPSRMAVMGH<br>SMGGGGALHAAANDTSLKAAIPLVGW<br>YPQTDWSSVKTPTLVVGAELDTIAPV<br>ETHSLAFYNSLPSDLDKAYLVLRGAS<br>HLVSTSPNTTIKKYTVAWLKRFVDND<br>RRYEKFLCPAPSDPRISEYRSTCPFL | 80.69 |

| | | | | | |
|---|---|---|---|---|---|
| E12 | PET selected filter e4 30 | 8 | LigandMPNN | MANPYERGPDPTESSIEAARGPFAVA<br>STTVSAAAADGFGGGTIYYPTDTSQG<br>RYGAVAISPGFTAGRSSIEWLGPAIA<br>SHGFVVITIDTITPLDQPDSRGDQLL<br>AALDYLRTHSAVKNRIDPSRMAVMGH<br>SMGGGGALYAAANDTSLKAAIPLVGW<br>HPRTDWSSVKTPTLVVGAELDTIAPV<br>ETHSRAFYDSLPADLDKAYLVLRGAS<br>HLVSTTPNTTILKYTVAWLRRFVDND<br>RRYEKFLCPAPSDPAISEYRSTCPFL | 81.08 |
| F1 | PET selected filter e4 30 | 9 | LigandMPNN | MANPYERGPDPTESSIEAERGPFAVA<br>STTVSAAEADGFGGGTIYYPTDTSQG<br>RYGAVAISPGFTAGRSSIEWLGPAIA<br>SHGFVVITIDTISLLDQPDSRGDQLL<br>AALDYLRTHSAVKNRIDPSRMAVMGH<br>SMGGGGALHAAAKDTSLKAAIPLVGW<br>HPQTDWSSVKTPTLVVGAELDTIAPV<br>ETHSKAFYDSLPSTLDKAYLELRGAS<br>HLVSTSPNRTILKYTVAWLRRFVDDD<br>RRYEKFLCPAPSDPAISEYRSTCPFL | 79.92 |
| F2 | PET selected filter e50 50 | 2 | LigandMPNN | MANPYVRGPDPTESSLSAASGPFAVA<br>QTTVSAAAAKGFGGGTIYYPTDTSQG<br>TYGAVAIAPGFTAGRESVAWWGPRIA<br>SQGFVVITIDTITLLDQPASRGRQLL<br>AALDYLRTNSVVKDKIDPSRMAVMGH<br>SMGGGGALHAAANDTSLEAAIPLQGW<br>DPRTDWSSVKTPTLVIGAQLDTIAPV<br>ASHSLAFYNSLPSTLNKAYLELAGAS<br>HLVSNTPDTTIARYTIAWLKRFVDRD<br>LRYEQFLCPAPSDPAISAYRSTCPFL | 83.01 |
| F3 | PET selected filter e50 50 | 3 | ProteinMPNN | MANPYIRGPDPTRSSLRARTGPFAVS<br>QTTVSREEAKGFGGGTIYYPTDTSQG<br>TYGAVAIAPGFTATRESVAWLGPRIA<br>SQGFVVITIDTISLLDQPESRGRQLL<br>AALDYLRTHSVVKDKIDPSRMAVMGH<br>SMGGGGALYAAAQDTSLEAAIPLQGW<br>SPQLDWSAVRTPTLVISAQLDTIAPP<br>ESHSLAFYNSLPADLNKAYLEIAGAS<br>HLVSNTWDDTIAEYTIAWLKRFVDRD<br>LRYDQFLCPAPSDPRISAYRSTCPFL | 78.38 |
| F4 | PET selected filter e50 70 | 3 | LigandMPNN | MANPYVKGPDPTEASLSAATGPFATA<br>STEVSAAAAKGFGGGTIYYPTDTSQG<br>RYGAVAIAPGFTAGRDSVEWLGPAIA<br>SHGFVVIVIDTITPTDGPATRGDQLL<br>AALDYLKTHSAVKDKIDPSRRGVMGH<br>SMGGGGALHAAANDSSLEAAIPLVGW<br>DPVTDWSSVKTPTLVIGAELDTIAPV<br>ETHSRAFYETLPSDLNKAYLELAGAS<br>HLVATSPNETILKYTVAWLKTFIDKD<br>KRYEKFLCPAPSDPAISAYDSTCPFK | 68.73 |

| | | | | | |
|---|---|---|---|---|---|
| F5 | PET selected format e10 30 | 8 | LigandMPNN | MANPYVKGPDPTEASISAATGPFAVA STTVSKAAAKGFGGGTIYYPTDTSQG TYGAVAIAPGFTAGRSSVEWLGPAIA SHGFVVITIDTITPTDGPATRGDQLL AALDYLKTNSAVKDKIDPSRMAVMGH SMGGGGALHAAANDTSLKAAIPLVGW DPRTDWSSVKTPTLFVGAELDTIAPV ATHTRAFYDSLPSTLNKAFLELRGAS HLVANTPDTTIKRYSIAWLRTFVDND RRYEQFLCPAPADPAISAYRSTCPFS | 74.52 |
| F6 | PET selected format e10 30 | 9 | LigandMPNN | MANPYVKGPDPTEASISADAGPFAVD SVTVSRAAAKGFGGGTIYYPTDTSQG TYGAVAIAPGFTAGRSSVEWLGPAIA SHGFVVITIDTITLTDGPATRGDQLL AALDYLKTNSAVKDKIDPSRMAVMGH SMGGGGALHAAANDTSLKAAIPLVGW DPRTDWSSVKTPTLIVGAELDTIAPV ATHSLAFYDSLPSDLNKAYLELRGAS HLVANTPDTTIKRYTIAWLRTFVDDD RRYEKFLCPAPSDPAISAYRSTCPFK | 74.9 |
| F7 | PET selected format e10 50 | 8 | LigandMPNN | MANPYVKGPEPTEASLSAAAGPFAVD STEVSAAEAKGFGGGTIYYPTDTSQG TYGAVAIAPGFTAGRSSVEWWGPAIA SHGFVVIVIDTTTLTDGPATRGDQLL AALDYLDTHPEVKDKIDPSRRAVMGH SMGGGGALHAAANDTSLKAAIPLVGW DPRTDWSSIKTPTLVIGAELDTIAPV ETHSRAFYNTLPSTLNKAYLELAGAS HLVATTPNTTILRYTVAWLRTFVDDD KRYEKFLCPAPSDPAISAYDSTCPFS | 68.73 |
| F8 | PET selected format e30 30 | 7 | LigandMPNN | MANPYVKGPDPTEASLAAARGPFAVA STTVSKAAAKGFGGGTIYYPTDTSQG TYGAVAIAPGFTAGRDSVEWLGPAIA SHGFVVITIDTITPLDEPATRGDQLL AALDYLKTNSVVKNKIDPSRMAVMGH SMGGGGALHAAANDTSLKAAIPLVGW DPRTDWSSVKTPTLVVGAELDTIAPV ATHSRAFYDTLPSDLDKAYLELRGAS HLVATTPNETILRYTVAWLRRFVDDD RRYERFLCPAPSDPAISAYRSTCPFS | 75.68 |
| F9 | PET selected format e4 30 | 1 | LigandMPNN | MANPYVRGPDPTEASIEAAAGPFAVD STTVSALEADGFGGGTIYYPTDTSQG RYGAVAISPGFTAGASSVAWWGPAIA SQGFVVIVIDTITPLDGPATRGDQLL AALDWLKTHPAVKDRIDPSRMAVMGH SMGGGGALHAAANNSSLKAAIPLVGW DPQTDWSSVKTPTLIVGAELDTIAPV ETHSLAFYNSLPSDLDKAYLELRGAS HLVPTTPNTTILRYSVAWLRRFVDDD LRYEKFLCPAPSDPRISAYRSTCPFK | 76.45 |

| | | | | | |
|---|---|---|---|---|---|
| F10 | PET selected format e4 30 | 9 | LigandMPNN | MANPYVRGPDPTEASIEAASGPFAVD<br>SVEVSRLEADGFGGGTIYYPTDTSQG<br>RYGAVAISPGFTAGRDSVEWLGPAIA<br>SKGFVVIVIDTITPLDKPAVRGDQLL<br>AALDYLKTHPAVKDRIDPSRMAVMGH<br>SMGGGGALHAAANNSSLKAAIPLVGW<br>DPRTDWSSVKTPTLIVGAELDTIAPV<br>ATHSRAFYDSLPSDLDKAYLELRGAS<br>HLVPTSPNTTILKYSVAWLRRFVDDD<br>LRYEKFLCPAPKDPRISAYRSTCPFS | 75.68 |
| F11 | PET selected format e50 30 | 8 | LigandMPNN | MANPYVKGPAPTESSLSARSGPFAVD<br>STTVTAAEAKGFGGGTIYYPTDTSQG<br>TYGAVAIAPGFTAGRDSVEWLGPAIA<br>SHGFVVIVIDTTTLTDGPATRGDQLL<br>AALDYLKTNPVVKNKIDPSRMAVMGH<br>SMGGGGALHAAANDTSLKAAIPLVGW<br>DPQTDWSSVKTPTLIVGAELDTIAPV<br>ETHSRAFYNSLPSTLDKAYLELRGAS<br>HLVATTPNETILKYSVAWLRRFVNDD<br>RRYEQFLCPAPSDPAISAYDSTCPFL | 73.36 |
| F12 | PET selected format e50 30 | 10 | LigandMPNN | MPNPYVKGPAPTESSLSAASGPFAVA<br>STTVTAAEAKGFGGGTIYYPTDTSQG<br>TYGAVAIAPGFTAGASSVAWLGPAIA<br>SKGFVVIVIDTTTLTVGPAERGDQLL<br>AALDYLKTNPVVKNKIDPSRMAVMGH<br>SMGGGGALHAAANDTSLKAAIPLVGW<br>DPQTDWSSVKTPTLIVGAELDTIAPV<br>ATHSRAFYDSLPSTLDKAYLELRGAS<br>HLVATTPNTTILRYTVAWLRRFVDND<br>RRYEQFLCPAPSDPAISAYDSTCPFL | 72.59 |

**Table S3. Rosetta Score (in kcal/mol) for PHL7 and specific residue substitutions in active PHL7 redesigns D5 and D11.** Residue substitutions in the redesigned variants with respect to PHL7 are highlighted in bold. Identical substitutions in both D5 and D11 are shown in pink. The last lines correspond to the per-residue energy values of the catalytic triad.

| Residue Position | PHL7 Residue | PHL7 Energy | D5 Residue | D5 Energy | D5 ΔEnergy | D11 Residue | D11 Energy | D11 ΔEnergy |
|---|---|---|---|---|---|---|---|---|
| 6 | GLU | -2.442 | GLU | -0.299 | 2.143 | **VAL** | **0.067** | **2.509** |
| 16 | ILE | -2.504 | **LEU** | **-4.701** | **-2.196** | **LEU** | **-3.997** | **-1.493** |
| 17 | GLU | -3.174 | **SER** | **-0.964** | **2.210** | **SER** | **-0.863** | **2.311** |
| 19 | VAL | 0.665 | VAL | 2.056 | 1.391 | **ASP** | **-2.103** | **-2.768** |
| 20 | ARG | -1.087 | **SER** | **-1.578** | **-0.491** | ARG | -0.492 | 0.595 |
| 27 | GLN | -0.638 | **SER** | **-1.609** | **-0.971** | **SER** | **-1.656** | **-1.018** |
| 33 | LEU | 2.477 | LEU | -0.016 | -2.493 | **ALA** | **1.060** | **-1.416** |
| 34 | GLN | -2.085 | **GLU** | **-1.775** | **0.310** | **GLU** | **-2.946** | **-0.861** |
| 36 | ASP | -0.482 | **LYS** | **0.523** | **1.005** | **LYS** | **0.573** | **1.055** |
| 54 | PHE | -1.763 | PHE | -2.948 | -1.185 | **TYR** | **-4.032** | **-2.269** |
| 60 | SER | -2.503 | SER | -2.250 | 0.252 | **ALA** | **-2.308** | **0.195** |
| 67 | GLN | -3.839 | **ARG** | **-3.529** | **0.310** | **ARG** | **-3.656** | **0.183** |
| 68 | GLU | -0.365 | **SER** | **-0.814** | **-0.449** | GLU | -1.154 | -0.789 |
| 70 | ILE | -4.237 | ILE | -1.679 | 2.558 | **VAL** | **-2.752** | **1.485** |
| 92 | ARG | -0.839 | **PRO** | **-4.128** | **-3.289** | **PRO** | **-4.088** | **-3.249** |
| 93 | LEU | -0.863 | **THR** | **-0.314** | **0.549** | LEU | -0.525 | 0.338 |
| 95 | GLN | -2.138 | **LEU** | **-0.423** | **1.715** | GLN | -2.077 | 0.061 |
| 97 | ASP | -1.280 | **ALA** | **0.861** | **2.140** | ASP | -1.048 | 0.232 |
| 98 | SER | -0.846 | **THR** | **-0.235** | **0.610** | SER | -1.042 | -0.197 |
| 101 | ARG | 0.000 | **ASP** | **-1.992** | **-1.992** | **ASP** | **-2.817** | **-2.817** |
| 104 | GLN | -1.905 | **LEU** | **-3.370** | **-1.466** | **LEU** | **-3.085** | **-1.180** |
| 109 | HIS | -4.320 | HIS | -3.943 | 0.377 | **TYR** | **-7.234** | **-2.914** |
| 113 | ASN | -4.121 | ASN | -1.009 | 3.112 | **HIS** | **-0.181** | **3.940** |
| 118 | ASN | -0.689 | ASN | -0.513 | 0.176 | **ASP** | **-1.320** | **-0.630** |
| 119 | ARG | -3.622 | ARG | -3.531 | 0.091 | **LYS** | **-3.809** | **-0.187** |
| 123 | ASN | -1.318 | **SER** | **-2.262** | **-0.945** | **SER** | **-2.364** | **-1.047** |
| 139 | SER | -1.051 | **HIS** | **-3.081** | **-2.030** | **HIS** | **-2.822** | **-1.771** |
| 144 | ASN | -3.333 | ASN | -2.606 | 0.727 | **ASP** | **-3.817** | **-0.483** |

| | | | | | | | | |
|---|---|---|---|---|---|---|---|---|
| 157 | HIS | -2.940 | **ASP** | **-4.277** | **-1.336** | HIS | -2.728 | 0.213 |
| 158 | THR | 0.579 | THR | -0.889 | -1.468 | **PRO** | **-1.393** | **-1.972** |
| 159 | ARG | -3.348 | **GLN** | **-3.257** | **0.091** | **GLN** | **-1.342** | **2.005** |
| 160 | LYS | -2.069 | **THR** | **0.468** | **2.537** | **THR** | **-1.579** | **0.490** |
| 161 | ASN | -2.792 | **ASP** | **-3.870** | **-1.078** | **ASP** | **-3.737** | **-0.946** |
| 166 | ARG | -0.803 | **LYS** | **-2.421** | **-1.618** | **LYS** | **-2.121** | **-1.317** |
| 172 | VAL | -5.046 | **ILE** | **-6.273** | **-1.226** | **ILE** | **-6.515** | **-1.469** |
| 175 | GLN | -1.349 | **GLU** | **-2.562** | **-1.213** | GLN | -2.084 | -0.735 |
| 183 | SER | -0.950 | SER | -0.851 | 0.098 | **PRO** | **-4.128** | **-3.289** |
| 184 | SER | -0.464 | **THR** | **-0.582** | **-0.118** | SER | -0.892 | -0.428 |
| 187 | GLU | -2.092 | **ARG** | **-2.209** | **-0.117** | **LEU** | **-1.755** | **0.337** |
| 191 | ASN | -0.325 | **ASP** | **-2.832** | **-2.507** | ASN | -1.291 | -0.967 |
| 192 | SER | -3.348 | **THR** | **-2.119** | **1.229** | SER | -1.253 | 2.096 |
| 196 | ASP | -0.353 | ASP | -0.681 | -0.328 | **THR** | **0.377** | **0.730** |
| 198 | ASP | -1.193 | ASP | -3.140 | -1.947 | **ASN** | **-1.933** | **-0.740** |
| 201 | TYR | -4.366 | **PHE** | **-3.426** | **0.940** | TYR | -5.391 | -1.025 |
| 202 | MET | -4.600 | **LEU** | **-5.375** | **-0.776** | **LEU** | **-4.713** | **-0.114** |
| 203 | GLU | -2.415 | **VAL** | **-5.859** | **-3.444** | GLU | -4.377 | -1.962 |
| 205 | ARG | -2.490 | **LYS** | **-1.181** | **1.309** | **ALA** | **-0.416** | **2.074** |
| 213 | ASN | -0.860 | **THR** | **-2.797** | **-1.938** | ASN | -1.936 | -1.076 |
| 219 | THR | -4.788 | **ILE** | **-6.561** | **-1.773** | **ILE** | **-6.835** | **-2.047** |
| 223 | SER | -2.780 | SER | -3.226 | -0.447 | **THR** | **-4.739** | **-1.959** |
| 233 | ASP | -1.147 | ASP | -0.897 | 0.249 | **LYS** | **-1.557** | **-0.410** |
| 235 | LEU | 0.705 | **ARG** | **-1.081** | **-1.786** | LEU | 0.569 | -0.137 |
| 238 | GLU | -2.156 | **ASP** | **-5.269** | **-3.112** | GLU | -3.530 | -1.374 |
| 239 | GLN | -1.491 | **LYS** | **-1.803** | **-0.312** | GLN | -1.312 | 0.179 |
| 246 | ASP | -0.127 | **SER** | **0.999** | **1.126** | **SER** | **-0.242** | **-0.114** |
| 248 | PHE | 2.659 | **PRO** | **-2.017** | **-4.676** | **PRO** | **-1.517** | **-4.176** |
| 252 | GLU | -2.739 | GLU | -3.296 | -0.557 | **ALA** | **-1.476** | **1.263** |
| 131 | SER | -2.499 | SER | -2.640 | -0.141 | SER | -2.900 | -0.401 |
| 177 | ASP | -4.948 | ASP | -5.942 | -0.994 | ASP | -5.835 | -0.888 |
| 209 | HIS | -3.497 | HIS | -3.446 | 0.051 | HIS | -3.060 | 0.437 |

**Table S4. Total Rosetta energy of the relaxed structures of PHL7 and native redesigns D5 and D11.**

| Structure | Total Energy (kcal/mol) | Δ(Total Energy) vs PHL7 |
|---|---|---|
| PHL7 | -804.161 | 0 |
| D5 | -839.738 | -35.577 |
| D11 | -843.765 | -39.604 |

**Table S5. Largest changes in per-residue energy (in kcal/mol) for D5 and D11 variants**

1. **10 best positive changes for D5**

| Residue Position | PHL7 Residue | PHL7 Energy | D5 Residue | D5 Energy | D5 ΔEnergy |
|---|---|---|---|---|---|
| 248 | PHE | 2.659 | PRO | -2.017 | -4.676 |
| 203 | GLU | -2.415 | VAL | -5.859 | -3.444 |
| 92 | ARG | -0.839 | PRO | -4.128 | -3.289 |
| 238 | GLU | -2.156 | ASP | -5.269 | -3.112 |
| 124 | ARG | -1.569 | ARG | -4.509 | -2.940 |
| 191 | ASN | -0.325 | ASP | -2.832 | -2.507 |
| 33 | LEU | 2.477 | LEU | -0.016 | -2.493 |
| 82 | PHE | -0.797 | PHE | -3.236 | -2.439 |
| 121 | ASP | -3.989 | ASP | -6.428 | -2.439 |
| 140 | ALA | -2.200 | ALA | -4.511 | -2.311 |

2. **10 best negative changes for D5**

| Residue Position | PHL7 Residue | PHL7 Energy | D5 Residue | D5 Energy | D5 ΔEnergy |
|---|---|---|---|---|---|
| 115 | VAL | -1.820 | VAL | 1.825 | 3.645 |
| 113 | ASN | -4.121 | ASN | -1.009 | 3.112 |
| 70 | ILE | -4.237 | ILE | -1.679 | 2.558 |
| 160 | LYS | -2.069 | THR | 0.468 | 2.537 |
| 25 | VAL | -3.399 | VAL | -1.032 | 2.367 |
| 17 | GLU | -3.174 | SER | -0.964 | 2.210 |
| 8 | GLY | -3.250 | GLY | -1.071 | 2.179 |
| 6 | GLU | -2.442 | GLU | -0.299 | 2.143 |
| 97 | ASP | -1.280 | ALA | 0.861 | 2.140 |
| 230 | PHE | -3.586 | PHE | -1.867 | 1.719 |

3. **10 best positive changes for D11**

| Residue Position | PHL7 Residue | PHL7 Energy | D11 Residue | D11 Energy | D11 ΔEnergy |
|---|---|---|---|---|---|
| 248 | PHE | 2.659 | PRO | -1.517 | -4.176 |
| 92 | ARG | -0.839 | PRO | -4.088 | -3.249 |
| 109 | HIS | -4.320 | TYR | -7.234 | -2.914 |
| 101 | ARG | 0.000 | ASP | -2.817 | -2.817 |
| 19 | VAL | 0.665 | ASP | -2.103 | -2.768 |
| 82 | PHE | -0.797 | PHE | -3.140 | -2.343 |
| 54 | PHE | -1.763 | TYR | -4.032 | -2.269 |
| 121 | ASP | -3.989 | ASP | -6.067 | -2.078 |
| 219 | THR | -4.788 | ILE | -6.835 | -2.047 |
| 114 | SER | -0.570 | SER | -2.604 | -2.035 |

### 4. 10 best negative changes for D11

| Residue Position | PHL7 Residue | PHL7 Energy | D11 Residue | D11 Energy | D11 ΔEnergy |
|---|---|---|---|---|---|
| 115 | VAL | -1.820 | VAL | 2.211 | 4.031 |
| 113 | ASN | -4.121 | HIS | -0.181 | 3.940 |
| 162 | TRP | -6.411 | TRP | -2.785 | 3.626 |
| 143 | ASN | -1.814 | ASN | 1.187 | 3.002 |
| 51 | GLN | -2.888 | GLN | -0.325 | 2.563 |
| 6 | GLU | -2.442 | VAL | 0.067 | 2.509 |
| 17 | GLU | -3.174 | SER | -0.863 | 2.311 |
| 8 | GLY | -3.250 | GLY | -1.140 | 2.110 |
| 122 | PRO | -3.913 | PRO | -1.816 | 2.097 |
| 192 | SER | -3.348 | SER | -1.253 | 2.096 |

**Table S6. Changes in salt bridges between the structures of PHL7, D5 and D11.** Residue pairs forming salt bridges in PHL7, D5, and D11, with their corresponding distances. Complex salt bridges where two residues simultaneously interact with another residue are highlighted in pink.

| PHL7 | D5 | D11 |
|---|---|---|
| | ARG67(C1) ↔ ASP88(C1) - 2.71 Å | ARG67(C1) ↔ ASP88(C1) - 2.84 Å |
| | | ARG76(C1) ↔ ASP19(C1) - 3.07 Å |
| ARG111(C1) ↔ ASP108(C1) - 3.27 Å | ARG111(C1) ↔ ASP108(C1) - 2.91 Å | ARG111(C1) ↔ ASP108(C1) - 2.65 Å |
| ARG124(C1) ↔ GLU148(C1) - 2.9 Å | ARG124(C1) ↔ ASP233(C1) - 2.77 Å | |
| | HIS139(C1) ↔ ASP157(C1) - 2.74 Å | HIS139(C1) ↔ ASP101(C1) - 2.84 Å |
| HIS157(C1) ↔ ASP97(C1) - 2.92 Å | | HIS157(C1) ↔ ASP97(C1) - 2.84 Å |
| | ARG187(C1) ↔ GLU252(C1) - 2.97 Å | |
| HIS209(C1) ↔ ASP177(C1) - 2.69 Å | HIS209(C1) ↔ ASP177(C1) - 2.74 Å | HIS209(C1) ↔ ASP177(C1) - 2.74 Å |
| LYS221(C1) ↔ GLU13(C1) - 2.69 Å | LYS221(C1) ↔ GLU13(C1) - 3.57 Å | LYS221(C1) ↔ GLU13(C1) - 2.9 Å |
| LYS221(C1) ↔ GLU17(C1) - 2.69 Å | LYS221(C1) ↔ ASP247(C1) - 2.91 Å | |
| LYS228(C1) ↔ ASP232(C1) - 2.97 Å | LYS228(C1) ↔ ASP232(C1) - 2.92 Å | LYS228(C1) ↔ ASP232(C1) - 2.89 Å |
| ARG229(C1) ↔ GLU238(C1) - 3.07 Å | | ARG229(C1) ↔ GLU238(C1) - 2.99 Å |
| ARG229(C1) ↔ ASP233(C1) - 3.52 Å | | |
| | | LYS233(C1) ↔ GLU148(C1) - 2.49 Å |
| | ARG235(C1) ↔ ASP238(C1) - 3.5 Å | |
| ARG236(C1) ↔ ASP10(C1) - 2.84 Å | ARG236(C1) ↔ ASP10(C1) - 3.15 Å | ARG236(C1) ↔ ASP10(C1) - 3.14 Å |
| ARG236(C1) ↔ ASP234(C1) - 2.88 Å | ARG236(C1) ↔ ASP234(C1) - 3.02 Å | ARG236(C1) ↔ ASP234(C1) - 2.99 Å |
| ARG254(C1) ↔ GLU252(C1) - 2.72 Å | | |
| ARG254(C1) ↔ GLU187(C1) - 2.94 Å | ARG254(C1) ↔ ASP191(C1) - 2.86 Å | |
